# Structural basis for enhanced infectivity and immune evasion of SARS-CoV-2 variants

**DOI:** 10.1101/2021.04.13.439709

**Authors:** Yongfei Cai, Jun Zhang, Tianshu Xiao, Christy L. Lavine, Shaun Rawson, Hanqin Peng, Haisun Zhu, Krishna Anand, Pei Tong, Avneesh Gautam, Shen Lu, Sarah M. Sterling, Richard M. Walsh, Sophia Rits-Volloch, Jianming Lu, Duane R. Wesemann, Wei Yang, Michael S. Seaman, Bing Chen

## Abstract

Several fast-spreading variants of severe acute respiratory syndrome coronavirus 2 (SARS-CoV-2) have become the dominant circulating strains that continue to fuel the COVID-19 pandemic despite intensive vaccination efforts throughout the world. We report here cryo-EM structures of the full-length spike (S) trimers of the B.1.1.7 and B.1.351 variants, as well as their biochemical and antigenic properties. Mutations in the B.1.1.7 protein increase the accessibility of its receptor binding domain and also the binding affinity for receptor angiotensin-converting enzyme 2 (ACE2). The enhanced receptor engagement can account for the increased transmissibility and risk of mortality as the variant may begin to infect efficiently infect additional cell types expressing low levels of ACE2. The B.1.351 variant has evolved to reshape antigenic surfaces of the major neutralizing sites on the S protein, rendering complete resistance to some potent neutralizing antibodies. These findings provide structural details on how the wide spread of SARS-CoV-2 enables rapid evolution to enhance viral fitness and immune evasion. They may guide intervention strategies to control the pandemic.

## Introduction

The COVID-19 pandemic, caused by severe acute respiratory syndrome coronavirus 2 (SARS-CoV-2) (*1*), has led to loss of human life in millions, as well as devastating socioeconomic disruptions worldwide. Although the mutation rate of the coronavirus is relatively low because of the proofreading activity of its replication machinery (*2*),several variants of concern (VOCs) have emerged in different countries, including the B.1.1.7 lineage first identified in the United Kingdom, the B.1.351 lineage in South Africa and the B.1.1.28 lineage in Brazil within a period of several months (*3–5*). These variants not only appear to spread more rapidly than the virus from the initial outbreak (i.e., the strain Wuhan-Hu-1; ref(*1*)), but also are more resistant to immunity elicited by the Wuhan-Hu-1 strain in the setting of either natural infection or vaccination (*6–8*). The B.1.1.7 variant is of particularly concern because it has been reported to be significantly more deadly (*9, 10*). Thus, understanding the underlying mechanisms of the increased transmissibility, risk of mortality and immune resistance of the new variants may facilitate development of intervention strategies to control the crisis.

SARS-CoV-2 is an enveloped positive-stranded RNA virus that depends on fusion of viral and target cell membranes to enter a host cell. This first key step of infection is catalyzed by the virus-encoded trimeric spike (S) protein, which is also a major surface antigen and thus an important target for development of diagnostics, vaccines and therapeutics. The S protein is synthesized as a single-chain precursor and subsequently cleaved by a furin-like protease into the receptor-binding fragment S1 and the fusion fragment S2 (Fig. S1; ref(*11*)). It is generally believed that binding of the viral receptor angiotensin converting enzyme 2 (ACE2) on the host cell surface to the receptor-binding domain (RBD) of S1, together with a second proteolytic cleavage by a cellular serine protease TMPRSS2 or endosomal cysteine proteases cathepsins B and L (CatB/L) in S2 (S2’ site; Fig. S1; ref(*12*)), induce large conformational changes and prompt dissociation of S1 and irreversible refolding of S2 into a postfusion structure, ultimately leading to membrane fusion (*13, 14*). In the prefusion conformation, S1 folds into four domains - NTD (N-terminal domain), RBD, and two CTDs (C-terminal domains), and they wrap around the prefusion structure of S2. The RBD can adopt two distinct conformations –“up” for a receptor-accessible state and “down” for a receptor-inaccessible state (*15*). Remarkable progress in structural biology of the S protein has been achieved rapidly since the early stage of the pandemic, advancing our knowledge on the SARS-CoV-2 entry process considerably (*15–28*). We have previously identified two structural elements - the FPPR (fusion peptide proximal region) and 630 loop, which appear to modulate the overall stability of the S protein, as well as the RBD conformation and thus the receptor accessibility (*22, 28*).

The S protein is the centerpiece of almost all the first-generation COVID-19 vaccines, which were developed shortly after the initial outbreak using the sequence of the Wuhan-Hu-1 strain (*29, 30*). Several of them have completed phase 3 trials, showing impressive protective efficacy with minimal side effects (*31, 32*), and they have therefore received Emergency Use Authorization (EUA) by various regulatory agencies throughout the world. These vaccines appear to have somewhat lower efficacy against the B.1.351 variant than against its parental strain (*6–8, 33*). In addition, this variant became completely resistant to many convalescent serum samples from those recovered from COVID-19 (*8*). How to address genetic diversity has therefore become a high priority for developing next-generation vaccines. In this study, we have characterized the full-length S proteins from the B.1.1.7 and B.1.351 variants and determined their structures by cryo-electron microscopy (cryo-EM), providing a structural basis for understanding the molecular mechanisms of the enhanced infectivity, risk of mortality and immune evasion of these two variants.

## Results

### Biochemical and antigenic properties of the intact S proteins from the new variants

To characterize the full-length S proteins with the sequences derived from natural isolates of the B.1.1.7 (hCoV-19/England/MILK-C504CD/2020) and B.1.351 (hCoV-19/South Africa/KRISP-EC-MDSH925100/2020) variants (Fig. S1), we first transfected HEK293 cells with the respective expression constructs and compared their membrane fusion activities with those of the full-length S constructs of their parental strains (Wuhan-Hu-1: D614, and its early variant with D614G substitution: G614 (*34*)). All S proteins expressed at comparable levels, as judged by western blot (Fig. S2A). The cells expressing these S proteins fused efficiently with the cells transfected with a human ACE2 construct (Fig. S2B), indicating that they are fully functional as a membrane fusogen. Consistent with our previous findings (*22, 28*), the G614 and B.1.351 variant S constructs showed slightly higher fusion activity than those of the D614 and B.1.1.7 variant, but the differences diminished when the transfection level increased.

To produce the full-length S proteins, we added a C-terminal strep-tag to the B.1.1.7 and B.1.351 S (Fig. S3A), expressed and purified these proteins under the identical conditions established for producing the D614 and G614 S trimers (*22, 28*). The B.1.1.7 protein eluted in three distinct peaks representing the prefusion S trimer, the postfusion S2 trimer and the dissociated S1 monomer, respectively (*22*), consistent with Coomassie-stained SDS-PAGE analysis (Fig. S3B). Nonetheless, the prefusion trimer was still the predominant form, accounting for >70% of the total protein. Like the G614 trimer (*28*), B.1.351 protein eluted as a single major peak, corresponding to the prefusion S trimer (Fig. S3B), with no obvious peaks for dissociated S1 and S2. SDS-PAGE analysis showed that the peaks for the prefusion trimers contained primarily the cleaved S1/S2 complex for both the proteins with the cleavage level moderately higher for B.1.351 S than for B.1.1.7 S. These results indicate that the B.1.351 and G614 S proteins have almost identical biochemical properties, while the B.1.1.7 S trimer is slightly less stable.

To assess antigenic properties of the purified prefusion S trimers from the new variants, we first measured by bio-layer interferometry (BLI) their binding to recombinant soluble ACE2 and S-directed monoclonal antibodies isolated from COVID-19 convalescent individuals. These antibodies target various epitopic regions on the S trimer, as defined by clusters of competing antibodies and designated RBD-1, RBD-2, RBD-3, NTD-1, NTD-2 and S2 (Fig. S4A; ref(*35*)). All but the last two clusters contain neutralizing antibodies. The B.1.1.7 variant bound substantially more strongly to the receptor than did its G614 parent, regardless of the oligomeric state of ACE2 (Figs. 1 and S4B; Table S1). The B.1.351 protein had higher affinity for monomeric ACE2, but slightly lower affinity for dimeric ACE2, than did the G614 trimer. In both cases, affinity for ACE2 of the B.1.351 protein was significantly lower than that of the B.1.1.7 variant. These results suggest that the mutations in the RBD of the B.1.1.7 variant, N501Y in particular, enhance receptor recognition, while the additional mutations in the B.1.351 variant (i.e., K417N and E484K) reduce ACE2 affinity to a level close to that of the G614 protein. All selected monoclonal antibodies bound the G614 S with reasonable affinities, and the B.1.1.7 variant showed a similar pattern but with substantially stronger binding to almost all the antibodies (Figs. 1 and S4B; Table S1). In contrast, the B.1.351 variant completely lost binding to the two RBD-2 antibodies, 32B6 and 12A2, as well as to the two NTD-1 antibodies, 12C9 and 83B6, while the affinities for the rest of the antibodies were the same as those of the G614 trimer. An RBD-3 antibody, 63C7, which recognizes the RBD-up conformation, bound more strongly to the B.1.1.7 trimer than to the G614 and B.1.351 proteins, for which it had affinities nearly equal to each other. To rule out potential artifacts associated with the proteins purified in detergent, we tested by flow cytometry binding of these antibodies and of a designed trimeric ACE2 construct to the S proteins expressed on the cell surfaces. As shown in Fig. S5, the binding results with the membrane-bound S trimers are largely in agreement with the data measured by BLI. In particular, the B.1.1.7 S had a substantially higher affinity for the trimeric ACE2 than did the G614 and B.1.351 proteins.

**Figure 1.**
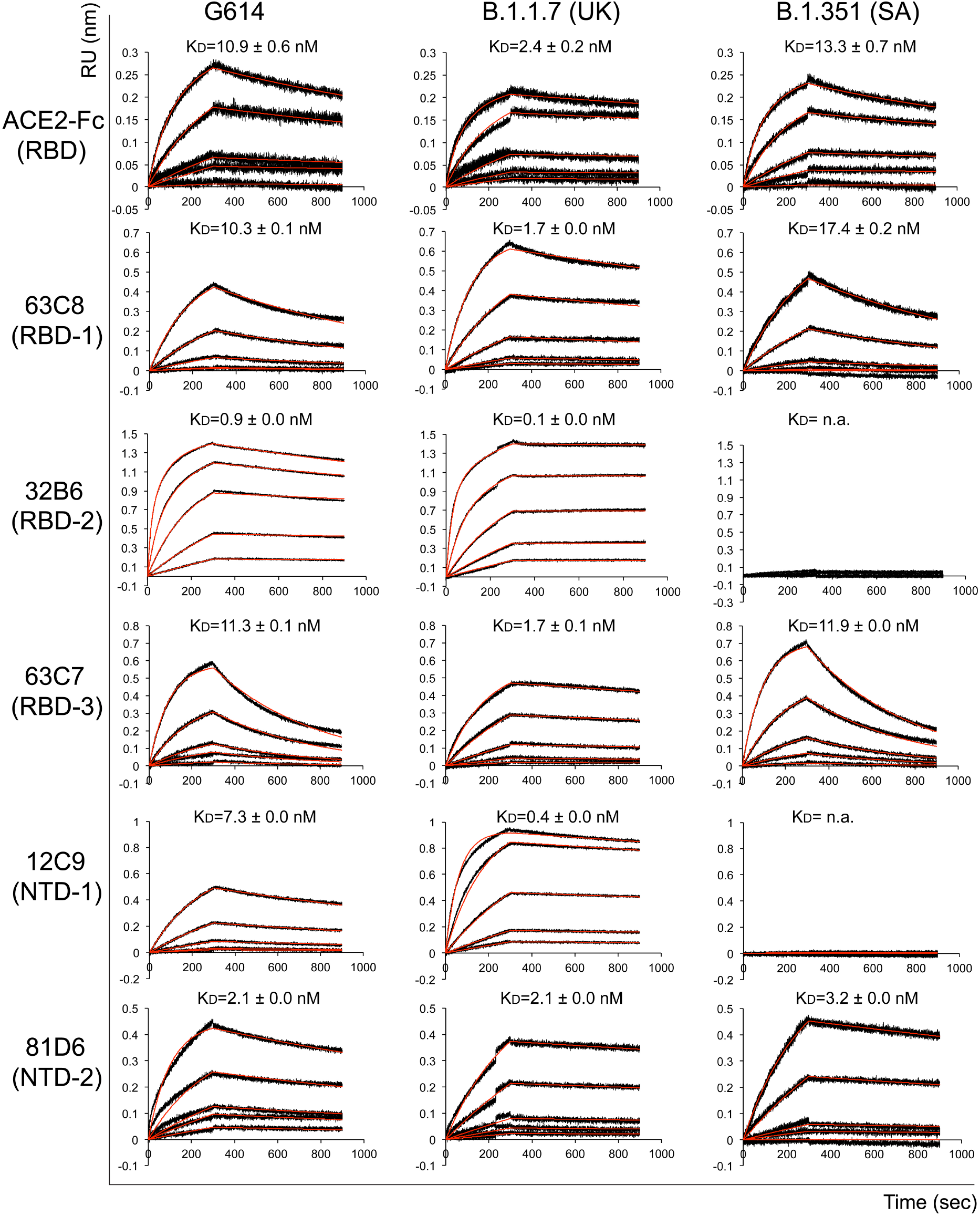
Antigenic properties of the purified full-length SARS-CoV-2 S proteins. Bio-layer interferometry (BLI) analysis of the association of prefusion S trimers from the G614 “parent” strain and the B.1.1.7 and B.1.351 variants derived from it with soluble ACE2 constructs and with a panel of antibodies representing five epitopic regions on the RBD and NTD (see Fig. S4A and ref(*35*)). For ACE2 binding, purified S proteins were immobilized to AR2G biosensors and dipped into the wells containing ACE2 at various concentrations. For antibody binding, various antibodies were immobilized to AHC biosensors and dipped into the wells containing each purified S protein at different concentration. Binding kinetics were evaluated using a 1:1 Langmuir model except for dimeric ACE2 and antibody 32B6 targeting the RBD-2, which were analyzed by a bivalent binding model. The sensorgrams are in black and the fits in red. RU, response unit. Binding constants are also summarized here and in Table S1. All experiments were repeated at least twice with essentially identical results.

We also assessed the neutralization potency of the antibodies and the trimeric ACE2 construct in blocking infection of these variants in an HIV-based pseudovirus assay. As summarized in Table S2, for most antibodies, the neutralization potency strictly correlated with their binding affinity for either the membrane-bound or purified S proteins. 81D6 and 163E6 recognize two non-neutralizing epitopes, located in the NTD and S2, respectively, and they did not neutralize any of the pseudoviruses. We note that the B.1.1.7 virus is the most sensitive to the trimeric ACE2 and the RBD-up-targeting 63C7, suggesting that the B.1.1.7 trimer may prefer the RBD-up conformation. These findings confirm that the detergent solubilized S proteins adopt a physiologically relevant conformation and demonstrate that mutations in the B.1.351 variant have a much greater impact on the antibody sensitivity of the virus than those in the B.1.1.7 variant.

### Structures of the full-length S trimers from the B.1.1.7 and B.1.351 variants

We determined the cryo-EM structures of the full-length S trimers with the unmodified sequences of the B.1.1.7 and B.1.351 variants. Cryo-EM images were acquired on a Titan Krios electron microscope operated at 300 keV and equipped with a Gatan K3 direct electron detector. We used RELION (*36*) and cryoSPARC (*37*) for particle picking, twodimensional (2D) classification, three dimensional (3D) classification and refinement (Figs. S6-S11). 3D classification identified five distinct classes for the B.1.1.7 S trimer --representing one closed prefusion conformation, three one-RBD-up conformations and one two-RBD-up conformation -- and two different classes for the B.1.351 trimer, representing a closed conformation and a one RBD-up conformation. These structures were refined to 3.1-4.5 Å resolution (Fig. S6-S9; Table S3).

As expected, the overall architectures of the full-length S proteins from the new variants are very similar to that of the G614 S trimer in the corresponding conformation (Figs. S10 and S11; ref(*28*)). In the closed, three RBD-down structure, the four domains of S1 -- NTD, RBD, CTD1 and CTD2 -- cap the almost invariant prefusion S2 trimer. In the one RBD-up conformation, the RBD position has no effect on the central core region of S2, but two NTDs, the immediately adjacent one and the one from the same protomer, shift away from the three-fold axis and open up the trimer. The furin cleavage site at the S1/S2 boundary in these structures remains disordered, and the structures therefore do not offer any explanation for the difference in the cleavage level between the B.1.1.7 and B.1.351 trimers; the position of a mutation (P681H) in the B.1.1.7 S (Fig. S1) close to the cleavage site is likewise not well ordered. A very small class of particles in the two RBD-up conformation was present only with the B.1.1.7 trimer (Fig. S10), possibly because of the greater number of uncleaved trimers from which S1 cannot dissociate.

For the B.1.1.7 S trimer, most of the particles used for refinement were in the RBD-up conformation (Fig. 2A-E). We have proposed that the FPPR and 630 loop modulate the stability and fusogenic structural rearrangements of the S protein (*22, 28*). In the closed conformation of the B.1.1.7 trimer, all three FPPR and three 630 loops are disordered (Fig. 2F), which otherwise would help clamp down the RBDs. This observation can explain why the B.1.1.7 trimer prefers an RBD-up conformation, unlike its parental stain, the G614 variant. In the one RBD-up conformation, one 630 loop on the opposite side of the shifted RBD becomes fully structured, inserting between neighboring NTD and CTDs in the same configuration found in the G614 trimer (*28*). The second 630 loop is partially ordered, while the third one remains disordered. A similar pattern is found for three FPPRs, although the structured FPPR adopts a conformation distinct from the one seen in our previous structures of the full-length S proteins (*22, 28*). Overall, the arrangement of these structural elements appears to stabilize the cleaved S trimer and to prevent the premature dissociation of S1 in the one RBD-up conformation. The three one RBD-up structures differ only by the degree to which the “up” RBD and the adjacent NTD of its neighboring protomer shift away from the central threefold axis (Fig. S12A). We have suggested that the two RBD-up conformation might be unstable (*22, 28*), leading to S1 dissociation and irreversible refolding of S2. If this suggestion is valid, the small class of the two RBD-up particles probably contains mainly uncleaved S trimers.

**Figure 2.**
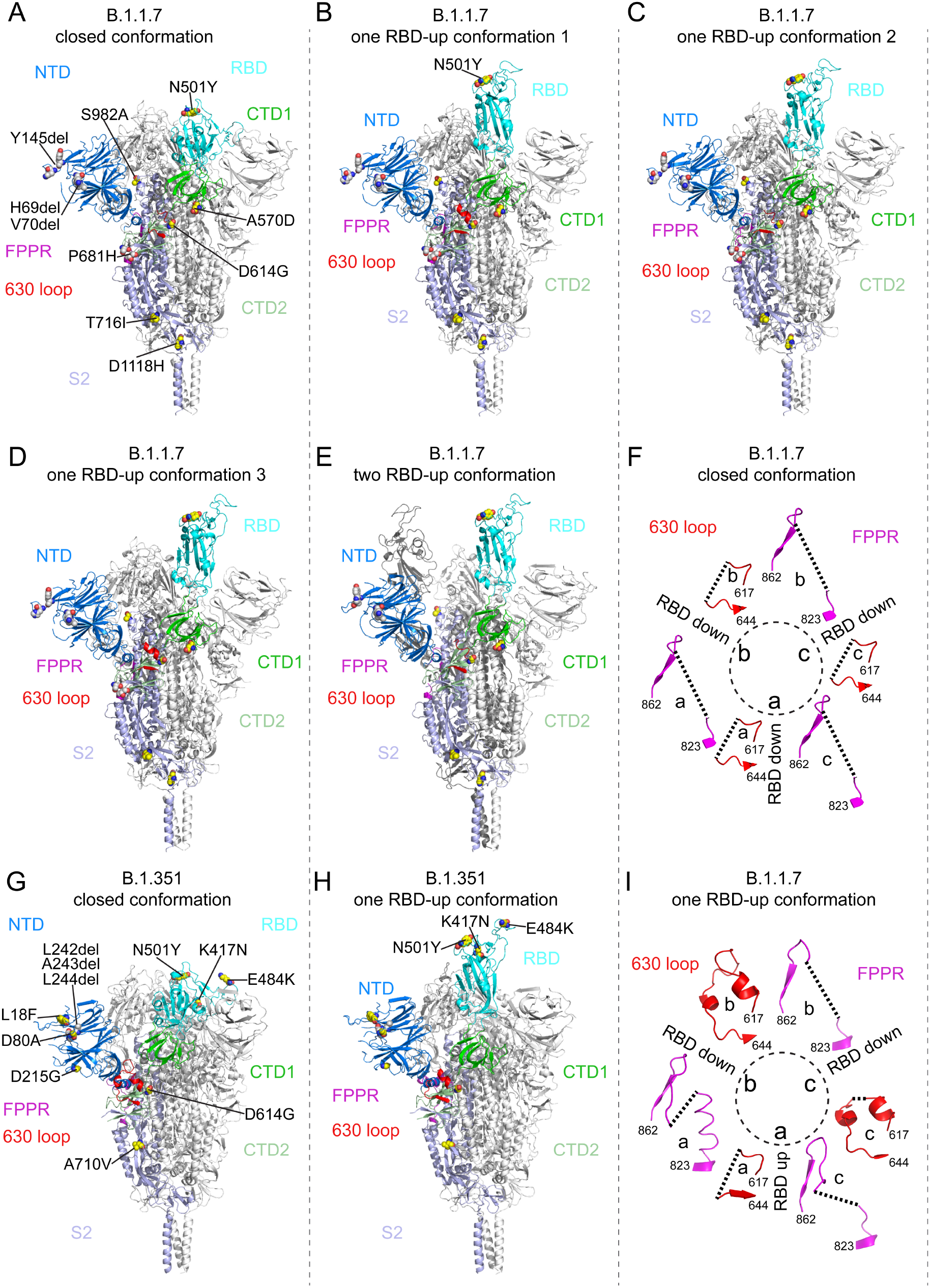
Cryo-EM structures of the full-length SARS-CoV-2 S proteins from the B.1.1.7 and B.1.351 variants. (**A-E**) The structures of the closed prefusion conformation, three one RBD-up conformations and a two RBD-up conformation of the B.1.1.7 S trimer are shown in ribbon diagram with one protomer colored as NTD in blue, RBD in cyan, CTD1 in green, CTD2 in light green, S2 in light blue, the 630 loop in red and the FPPR in magenta. (**G**) and (**H**) The structures of the closed prefusion conformation and one RBD-up conformation of the B.1.351 S trimer are shown in ribbon diagram with the same color scheme as in (A). All mutations in the new variants, as compared to the original virus (D614), are highlighted in sphere model. (**F**) and (**I**) Structures, in the B.1.1.7 trimer, of segments (residues 617-644) containing the 630 loop (red) and segments (residues 823-862) containing the FPPR (magenta) from each of the three protomers (a, b and c). The position of each RBD is indicated. Dashed lines indicate gaps in the chain trace (disordered loops).

The two classes for the B.1.351 S trimer represent the closed prefusion and one RBD-up states, respectively (Fig. 2G and 2H). The configurations of the FPPR and 630 loop follow closely the distribution seen in the G614 trimer: all are structured in the RBD-down conformation, while only one the FPPR and 630-loop pair is ordered in the one RBD-up conformation (Fig. S11; ref(*28*)). These observations are consistent with the nearly identical biochemical stabilities of the B.1.351 and G614 S trimers (Fig. S3; ref(*28*)).

### Structural consequences of mutations in the B.1.1.7 variant

We superposed the structures of the B.1.1.7 trimer onto the G614 trimer in the closed conformation aligning them by the S2 structure (Fig. 3A), and detected an outward rotation of all three S1 subunits, leading to a slightly more open conformation than that of the G614 trimer. This rotation in the B.1.1.7 variant widened the gap between the NTD and the CTDs of the same protomer (Fig. S12B). In the G614 trimer, this gap accommodates the ordered 630 loop that reinforces CTD2 and prevents S1 shedding (*28*). Thus, the widened gap in the variant loosens the grip on the 630 loop, accounting for the absence of ordered features in this part of the B.1.1.7 map. There are two mutations that may be responsible for these structural differences. First, Ala570 in CTD1 normally packs against one side of the FPPR in the G614 trimer (Fig. 3B). The A570D mutation, with a larger side chain, may weaken the packing and destabilize the FPPR, which is not visible in the closed conformation of the B.1.1.7 S. Moreover, in the one RBD-up conformation of the B.1.1.7 S, in which the FPPR is at least partially structured, Lys854, which in the G614 trimer probably forms a hydrogen bond with the main chain carbonyl group of Gly614, flips back in the B.1.1.7 variant to form a salt bridge with the mutantAsp570. Second, the S982A mutation eliminates a hydrogen bond between the central helices of S2 and the carbonyl group of Gly545 in CTD1 (Fig. 3C). These two mutations together allow an outward movement of CTD1 by more than 3Å (Fig. S12B), thereby affecting the conformation of both the FPPR and 630 loops.

**Figure 3.**
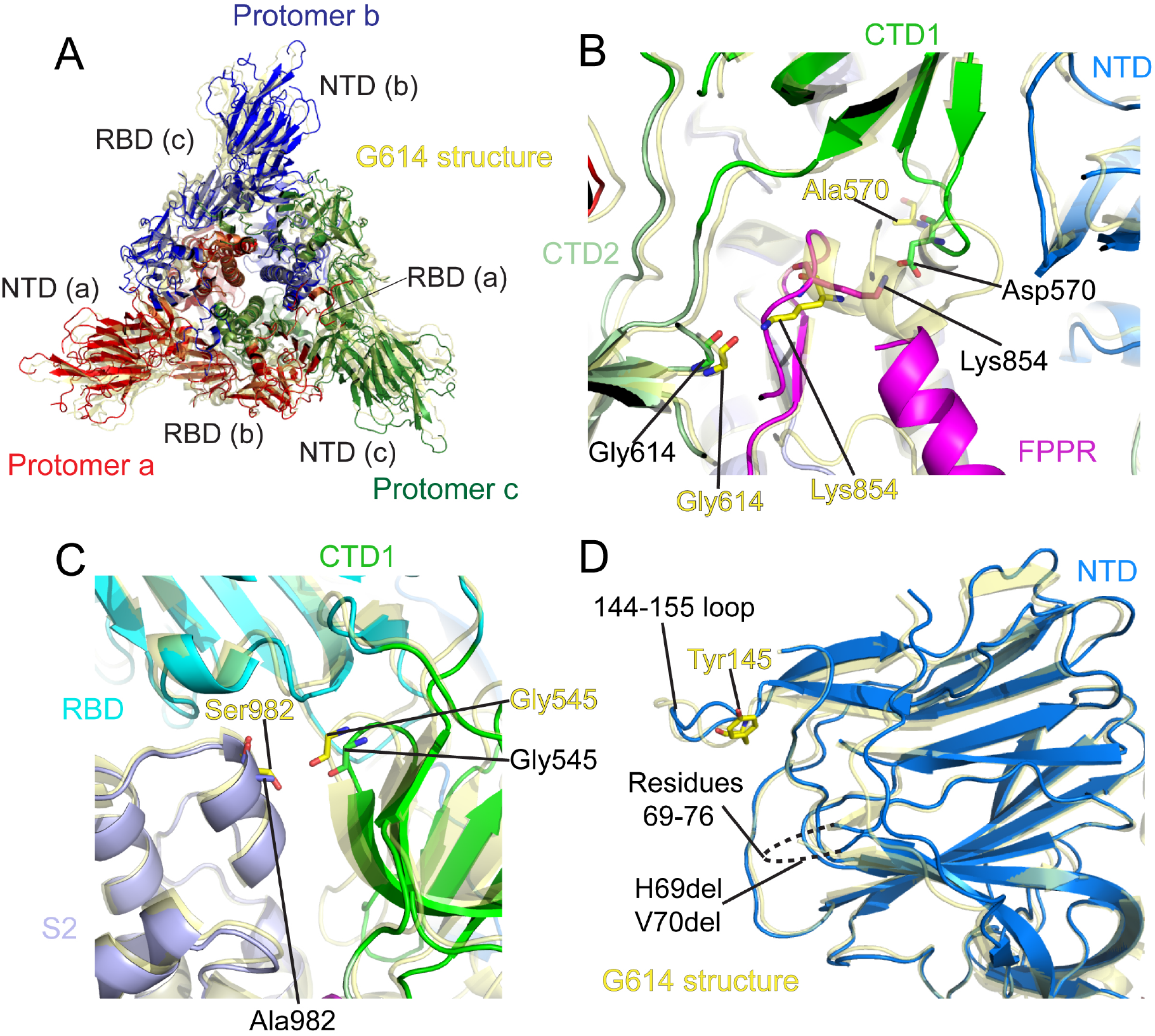
Structural impact of the mutations in the B.1.1.7 S. (**A**) Top views of superposition of the structure of the B.1.1.7 S trimer in ribbon representation with the structure of the prefusion trimer of the G614 S (PDB ID: 7KRQ), shown in yellow. NTD and RBD of each protomer are indicated. (**B**) A close-up view of the region near the A570D mutation with superposition of the B.1.1.7 trimer structure (one RBD-up) in green (CTD1) and magenta (FPPR) and the G614 trimer (closed) in yellow. Residues A570, D570, two G614 and two K854 from both structures are shown in stick model. (**C**) A view of the region near the S982A mutation with superposition of the B.1.1.7 trimer structure (closed) in green (CTD1) and magenta (FPPR) and the G614 trimer (closed) in yellow. (**D**) Superposition of the NTD structure of the B.1.1.7 S trimer in blue with the NTD of the G614 S trimer in yellow. Locations of Tyr145 and the disordered loop containing residues 69-76 are indicated.

Other mutations in the B.1.1.7 variant cluster in the NTD, including deletions of His69, Val70 and Tyr145 (Fig. 3D). The first two residues are in a disordered loop in all these S structures, and the structural impact, if any, of their deletion is unclear. Tyr145 is also near a loop (residues 144-155), and its deletion apparently causes only some local changes of the loop. The absence of structural changes in the B.1.1.7 trimer NTD is consistent with the absence of effects on its sensitivity to the various antibodies that target this domain (*35*). Additional mutations, including N501Y, T716I and D1118H, caused minimal local changes (Fig. S13A-C). They might nonetheless have important roles in the dynamic process of membrane fusion.

### Structural impact of the mutations in the B.1.351 variant

We compared the structures of the B.1.351 trimer with the G614 trimer in both the closed and the one RBD-up conformation (Figs. 4A and S14). The overall structures of variant and parent were essentially the same for the corresponding states, except for some loop regions in the NTD. Three mutations in the RBD, including K417N, E484K and N501Y, do not produce any major structural rearrangements (Fig. 4B). The most striking differences are in the NTD, which contains three point mutations (L18F, D80A and D215G) and a three-residue deletion (L242del, A243del and L244del). The L18F and D80A lead to reconfiguration of the N-terminal segment despite the disulfide between Cys16 and Cys136 that partly anchors the N-terminal peptide (Fig. 4C). D215G appears to have the least structural impact since Asp215 is a solvent-exposed residue that may compensate for the surface charge from the neighboring, well-exposed Arg214.

**Figure 4.**
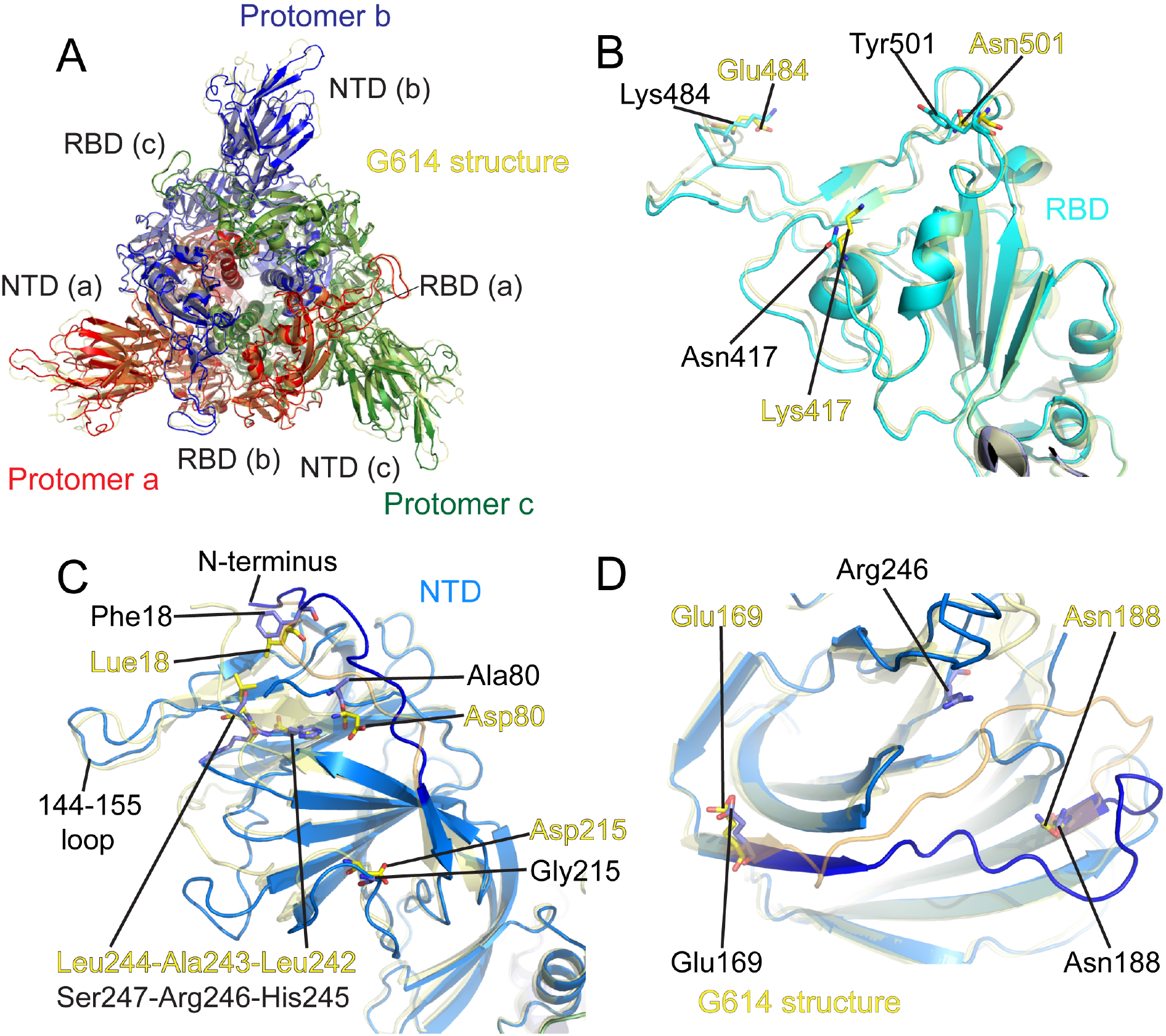
Structural impact of the mutations in the B.1.351 S. (**A**) Top views of superposition of the structure of the B.1.351 S trimer in ribbon representation with the structure of the prefusion trimer of the G614 S (PDB ID: 7KRQ), shown in yellow. NTD and RBD of each protomer are indicated. (**B**) Superposition of the RBD structure of the B.1.1.7 S trimer in blue with the RBD of the G614 S trimer in yellow. Locations of mutations K417N, E484K and N501Y are indicated and these residues are shown in stick model. (**C**) A view of the NTDs from superposition of the structure of the B.1.351 S trimer in blue and the G614 S in yellow. Locations of mutations L18F, D80A and D215G, as well as replacement of L242-A243-L244 by H245-R-246-A247 are indicated and the residues are shown in stick model. (**D**) Superposition of the NTD structure of the B.1.1.7 S trimer in blue with the NTD of the G614 S trimer in yellow. Displacement of the segment 169-188 and the location of R246 in the B.1.351 structure are indicated.

The most consequential changes are probably from the triple residue deletion, as these nonpolar residues, located on the edge of the NTD core structure formed by four stacking β-sheets, are replaced with polar residues His245-Arg246-Ser247. This replacement causes a shift of the nearby loop 144-155 and must also reconfigure the adjacent disorder loop (residues 246-260), both of which form part of the neutralizing epitopes (*38*). Furthermore, Arg246 is pointing towards the side chain of Arg102 near the segment 172-188, forcing the loop to rearrange. As shown in Fig. 4D, the segment 172-188 segment wraps around the edge of the NTD core, packing underneath the β-sheet involving L242-A243-L244 in the G614 trimer. The triple residue deletion rearranges this segment with a movement up to 17Å (Leu180), bringing large changes in the antigenic surface in this region. Thus, these mutations are likely to alter substantially the conformational preferences of this component of the molecular surface and to affect binding of any antibody that has part of its footprint in this region. Additional mutation A701V is located in surface-exposed region of S2 and caused minimal structural changes (Fig. S13D). It is difficult to predict the impact of this mutation since the functional role of this segment is unclear.

## Discussion

Transmissibility and immune evasion are independent selective forces driving emergence of viral genetic diversity. The changes of most concern in the SARS-CoV-2 S protein would be those that simultaneously enhance transmission, augment disease severity, and evade immune recognition in previously exposed hosts. Our data suggest that the most problematic combination of such mutations is not yet present in the existing variants examined here.

In the B.1.1.7 virus, mutations A570D and S982A lead to an outward shift of the CTD1 away from the threefold axis, thereby relaxing the two key structural elements, the FPPR and 630 loop, which in the parental strain help retain the RBD in its “down” position. The mutations effectively increase the frequency with which the S trimer samples the one RBD-up conformation and thus more effectively presents the receptor binding motif (RBM) to ACE2 on the host cells. Once one RBD flips up, the fully or partially ordered 630 loops of the neighboring protomers stabilize the CTD2, which folds together with the N-terminal segment of S2, and thus prevent the premature dissociation of S1 and inactivation of the functional spike. The N501Y mutation in the ACE2 binding site of the RBD also increases the affinity of that domain for both the monomeric and dimeric ACE2 proteins, probably because of hydrophobic interaction of Tyr501 with Tyr41 of ACE2 (*39*), as well as a possible cation-πinteraction with ACE2 Lys353 (Fig. S15). The combination of enhanced presentation of the RBM and additional local interactions might allow the B.1.1.7 virus to infect cell types with lower levels of ACE2 expression than those of the nasal and bronchial epithelial cells that the virus typically infects; an expanded cell tropism could account for the recently reported increased risk of mortality in patients infected with this variant (*9, 10*).

Residue 501 is outside of the major neutralizing epitope clusters in the RBD (Fig. S4), and its mutation from asparagine to tyrosine does not cause any widespread conformational shifts. The N501Y mutation has therefore brought only minimal changes in the sensitivity of the B.1.1.7 variant to the potently neutralizing antibodies that bind the RBD (Table S1 and S2). In addition, the mutations in the NTD caused no major structural rearrangements, consistent with potent neutralization against the B.1.1.7 virus by NTD-directed neutralizing antibodies. Thus, the B.1.1.7 variant, while posing a threat, as documented, to unvaccinated or virus-naïve individuals, may be a less serious concern for those with infection- or vaccine-elicited immunity to the Wu_Hu-1 strain (*33*).

In the B.1.351 virus, the S protein largely retains the structure of the G614 trimer with almost identical biochemical stability. Mutations N501Y, K417N and E484K in the RBD have not led to any major structural changes, but the loss of salt bridges between K417 and ACE2 Asp30 and Glu484 and ACE2 Lys31 more or less cancels the increased receptor affinity imparted by N501Y (Fig. S15). The mutations at residues 417 and 484 are probably responsible, however, for loss of detectable binding and neutralization by antibodies that target the RBD-2 epitopes (Fig. S4A). Moreover, the accompanying mutations in the NTD remodel the antigenic surface of the domain, greatly reducing the potency of neutralizing antibodies that target NTD-1 epitopes. In particular, deletion of three consecutive residues in a β-strand forming the core structure of the NTD is clearly an effective strategy to rearrange a large surface of the domain. The B.1.351 variant was probably selected under a certain level of immune pressure, as it altered two major neutralizing sites on the S trimer simultaneously with only a slight compromise in its ability to engage a host cell.

The global range of SARS-CoV-2 and the consequently vast number of replication events occurring every day make emergence of new variants inevitable, and substantially increases the genetic diversity of the virus. In many cases, antibody resistance may compromise viral fitness, as in the B.1.351 variant, which resists neutralization by RBD-directed antibodies, but also loses the enhanced affinity and transmissibility imparted by N501Y, as a consequence of the immune-escape mutations. It is also possible to combine immune evasion and virulence through continuous viral evolution, such as the identification of a B.1.1.7 variant that contains the E484K mutation (B.1.1.7+E484K) (*40*), although this added mutation may alleviate the risk of mortality, at least to some extent. Such a combination will nonetheless bring much greater challenges for vaccine development compared to the beginning of the pandemic. Genetic diversity is also the major hurdle for development or optimization of vaccines against several other human pathogens, such HIV-1, hepatitis C virus and influenza virus. If SARS-CoV-2 becomes seasonal, innovative strategies already developed against those viruses may be applicable to on-going control of the COVID-19 pandemic. The B.1.351 S trimer, which has superior biochemical stability and novel epitopes, should be an excellent starting point for developing next-generation vaccines designed to elicit broadly neutralizing antibody responses.

## Acknowledgments

We thank the SBGrid team for technical assistance, K. Arnett for support and advice on the BLI experiments, and S. Harrison and A. Carfi for critical reading of the manuscript. EM data were collected at the Harvard Cryo-EM Center for Structural Biology of Harvard Medical School. We acknowledge support for COVID-19 related structural biology research at Harvard from the Nancy Lurie Marks Family Foundation and the Massachusetts Consortium on Pathogen Readiness (MassCPR). This work was supported by NIH grants AI147884 (to B.C.), AI141002 (to B.C.), AI127193 (to B.C. and James Chou), a Fast grant by Emergent Ventures (to B.C.) and a COVID-19 Award by Massachusetts Consortium on Pathogen Readiness (MassCPR; to B.C.).

## Author Contribution

B.C., Y.C., J.Z. and T.X. conceived the project. Y.C. and H.P. expressed and purified the full-length S proteins. T.X. performed BLI and cell-cell fusion experiments. J.Z. and Y.C. prepared cryo grids and performed EM data collection with contributions from S.M.S. and R.M.W. J.Z. and Y.C. processed the cryo-EM data, built and refined the atomic models with help from S.R.. C.L.L. and M.S.S performed the neutralization assays using the HIV-based pseudoviruses. H.Z., K.A. and W.Y. performed the flow cytometry experiments. P.T., A.G. and, D.R.W. produced anti-S monoclonal antibodies. S.L. and J.L. created all the expression constructs. S.R.V. contributed to cell culture and protein production. All authors analyzed the data. B.C., Y.C., J.Z. and T.X. wrote the manuscript with input from all other authors.

## Materials and Methods

### Expression constructs

Genes of full-length spike (S) protein from hCoV-19/England/MILK-C504CD/2020 (GISAID accession ID: EPI_ISL_736724) and hCoV-19/South Africa/KRISP-EC-MDSH925100/2020 (GISAID accession ID: EPI_ISL_736980) were synthesized by GENEWIZ (South Plainfield, NJ). The S genes were fused with a C-terminal twin Strep tag [(GGGGS)2WSHPQFEK(GGGGS)_2_WSHPQFEK)] and cloned into a mammalian cell expression vector pCMV-IRES-puro (Codex BioSolutions, Inc, Gaithersburg, MD).

### Expression and purification of recombinant proteins

Expression and purification of the full-length S proteins were carried out as previously described (*22*). Briefly, expi293F cells (ThermoFisher Scientific, Waltham, MA) were transiently transfected with the S protein expression constructs. To purify the S protein, the transfected cells were lysed in a buffer containing Buffer A (100 mM Tris-HCl, pH 8.0, 150 mM NaCl, 1 mM EDTA) and 1% (w/v) n-dodecyl-β-D-maltopyranoside (DDM) (Anatrace, Inc. Maumee, OH), EDTA-free complete protease inhibitor cocktail (Roche, Basel, Switzerland), and incubated at 4°C for one hour. After a clarifying spin, the supernatant was loaded on a strep-tactin column equilibrated with the lysis buffer. The column was then washed with 50 column volumes of Buffer A and 0.3% DDM, followed by additional washes with 50 column volumes of Buffer A and 0.1% DDM, and with 50 column volumes of Buffer A and 0.02% DDM. The S protein was eluted by Buffer A containing 0.02% DDM and 5 mM desthiobiotin. The protein was further purified by gel filtration chromatography on a Superose 6 10/300 column (GE Healthcare, Chicago, IL) in a buffer containing 25 mM Tris-HCl, pH 7.5, 150 mM NaCl, 0.02% DDM.

The monomeric ACE2 or dimeric ACE2 proteins were produced as described (*41*). Briefly, Expi293F cells transfected with monomeric ACE2 or dimeric ACE2 expression construct and the supernatant of the cell culture was collected. The monomeric ACE2 protein was purified by affinity chromatography using Ni Sepharose excel (Cytiva Life Sciences, Marlborough, MA), followed by gel filtration chromatography. The dimeric ACE2 protein was purified by GammaBind Plus Sepharose beads (GE Healthcare), followed gel filtration chromatography on a Superdex 200 Increase 10/300 GL column. All the monoclonal antibodies were produced as described (*35*).

### Western blot

Western blot was performed using an anti-SARS-COV-2 S antibody following a protocol described previously (*42*). Briefly, full-length S protein samples were prepared from cell pellets and resolved in 4-15% Mini-Protean TGX gel (Bio-Rad, Hercules, CA) and transferred onto PVDF membranes. Membranes were blocked with 5% skimmed milk in PBS for 1 hour and incubated a SARS-CoV-2 (2019-nCoV) Spike RBD Antibody (Sino Biological Inc., Beijing, China, Cat: 40592-T62) for another hour at room temperature. Alkaline phosphatase conjugated anti-Rabbit IgG (1:5000) (Sigma-Aldrich, St. Louis, MO) was used as a secondary antibody. Proteins were visualized using one-step NBT/BCIP substrates (Promega, Madison, WI).

### Cell-cell fusion assay

The cell-cell fusion assay, based on the α-complementation of E. coli β-galactosidase, was conducted to quantify the fusion activity mediated by SARS-CoV2 S protein, as described (*22*). Briefly, various amount of the full-length SARS-CoV2 (D614, G614, UK and South Afirca) S construct (0.025-10 μg) and the α fragment of E. coli β-galactosidase construct (10 μg), or the full-length ACE2 construct (10 μg) together with the ω fragment of E. coli β-galactosidase construct (10 μg), were transfected to HEK293T cells using Polyethylenimine (PEI) (80 μg). After a 24-hour incubation at 37°C, the cells were detached using PBS buffer with 5mM EDTA and resuspended in complete DMEM medium. 50 μl S-expressing cells (1.0×10^6^ cells/ml) were mixed with 50 μl ACE2-expressing cells (1.0×10^6^ cells/ml) to allow the cell-cell fusion proceed at 37°C for 2 hours. Cell-cell fusion activity was quantified using a chemiluminescent assay system, Gal-Screen (Applied Biosystems, Foster City, CA), following the standard protocol recommended by the manufacturer. The substrate was added to the mixture of the cells and allowed to react for 90 minutes in dark at room temperature. The luminescence signal was recorded with a Synergy Neo plate reader (Biotek, Winooski, VT).

### Binding assay by bio-layer interferometry (BLI)

Binding of monomeric or dimeric ACE2 to the full-length Spike protein of each variant was measured using an Octet RED384 system (ForteBio, Fremont, CA), following the protocol described previously(*41*). Briefly, the full-length S protein was immobilized to Amine Reactive 2nd Generation (AR2G) biosensors (ForteBio, Fremont, CA) and dipped in the wells containing the ACE2 protein at various concentrations (5.56-450 nM for monomeric ACE2; 0.926-75 nM for dimeric ACE2) for 5 minutes, followed with 10 minutes in the running buffer (PBS, 0.02% Tween 20, 2 mg/ml BSA) to determine the dissociation rate. To measure the binding of the full-length Spike protein to monoclonal antibodies, the antibody was immobilized to anti-human IgG Fc Capture (AHC) biosensor (ForteBio, Fremont, CA) following a protocol recommended by the manufacturer. The full-length Spike protein was diluted using the running buffer (PBS, 0.02% Tween 20, 0.02% DDM, 2 mg/ml BSA) to various concentrations (0.617-50 nM) and transferred to 96-well plate. The sensors were dipped in the wells containing the Spike protein solutions for 5 minutes to measure the association rate, followed with 10 minutes in the wells of the running buffer to measure the dissociation rate. Control sensors with no Spike protein or antibody were also dipped in the ACE2 or Spike protein solutions and the running buffer as references. Recorded sensorgrams with background subtracted from the references were analyzed using the software Octet Data Analysis HT Version 11.1 (ForteBio). Binding kinetics was evaluated using a 1:1 Langmuir model except for dimeric ACE2 and antibodies 32B6 and 12A2, which were analyzed by a bivalent binding model.

### Flow cytometry

Expi293F cells (ThermoFisher Scientific) were grown in Expi293 expression medium (ThermoFisher Scientific). Cell surface display DNA constructs for the SARS-CoV-2 spike variants together with a plasmid expressing blue fluorescent protein (BFP) were transiently transfected into Expi293F cells using ExpiFectamine 293 reagent (ThermoFisher Scientific) per manufacturer’s instruction. Two days after transfection, the cells were stained with primary antibodies or the histagged ACE2_615_-foldon T27W protein (*41*) at 10 μg/ml concentration. For antibody staining, an Alexa Fluor 647 conjugated donkey anti-human IgG Fc F(ab’)2 fragment (Jackson ImmunoResearch, West Grove, PA) was used as secondary antibody at 5 μg/ml concentration. For ACE2_615_-foldon T27W staining, APC conjugated anti-HIS antibody (Miltenyi Biotec, Auburn, CA) was used as secondary antibody at 1:50 dilution. Cells were run through an Intellicyt iQue Screener Plus flow cytometer. Cells gated for positive BFP expression were analyzed for antibody and ACE2_615_-foldon T27W binding. The flow cytometry assays were repeated three times with essentially identical results.

### HIV-based pseudovirus assay

Neutralizing activity against SARS-CoV-2 pseudovirus was measured using a single-round infection assay in 293T/ACE2 target cells. Pseudotyped virus particles were produced in 293T/17 cells (ATCC) by co-transfection of plasmids encoding codon-optimized SARS-CoV-2 full-length Spike constructs, packaging plasmid pCMV DR8.2, and luciferase reporter plasmid pHR’ CMV-Luc. G614 Spike, packaging and luciferase plasmids were kindly provided by Dr. Barney Graham (Vaccine Research Center, NIH). The 293T cell line stably overexpressing the human ACE2 cell surface receptor protein was kindly provided by Drs. Michael Farzan and Huihui Ma (The Scripps Research Institute). For neutralization assays, serial dilutions of monoclonal antibodies (mAbs) were performed in duplicate followed by addition of pseudovirus. Pooled serum samples from convalescent COVID-19 patients or pre-pandemic normal healthy serum (NHS) were used as positive and negative controls, respectively. Plates were incubated for 1 hour at 37°C followed by addition of 293/ACE2 target cells (1×10^4^/well). Wells containing cells + pseudovirus (without sample) or cells alone acted as positive and negative infection controls, respectively. Assays were harvested on day 3 using Promega BrightGlo luciferase reagent and luminescence detected with a Promega GloMax luminometer. Titers are reported as the concentration of mAb that inhibited 50% or 80% virus infection (IC_50_ and IC_80_ titers, respectively). All neutralization experiments were repeated twice with similar results.

### Cryo-EM sample preparation and data collection

To prepare cryo grids, 3.5 μl of the freshly purified sample from the peak fraction in DDM at ~2.0 mg/ml for the B.1.1.7 protein or ~1.5 mg/ml for the B.1.351 protein was applied to a 1.2/1.3 Quantifoil grid (Quantifoil Micro Tools GmbH), which had been glow discharged with a PELCO easiGlow™ Glow Discharge Cleaning system (Ted Pella, Inc.) for 60 s at 15 mA. Grids were immediately plunge-frozen in liquid ethane using a Vitrobot Mark IV (ThermoFisher Scientific), and excess protein was blotted away by using grade 595 filter paper (Ted Pella, Inc.) with a blotting time of 4 s, a blotting force of −12 at 4°C in 100% humidity. For data collection, images were acquired with selected grids using a Titan Krios transmission electron microscope (ThermoFisher Scientific) operated at 300 keV and equipped with a BioQuantum GIF/K3 direct electron detector. Automated data collection was carried out using SerialEM version 3.8.6 (*43*) at a nominal magnification of 105,000× and the K3 detector in counting mode (calibrated pixel size, 0.825 Å) at an exposure rate of 20.21 (for B.1.1.7) or 20.50 (for B.1.351) electrons per pixel per second. Each movie had a total accumulated electron exposure of ~53.40 (for B.1.1.7) or 51.10 (for B.1.351) e/Å^2^, fractionated in 51 (for B.1.1.7) or 50 (for B.1.351) frames. Datasets were acquired using a defocus range of 1.4-2.3 μm (for B.1.1.7) or 1.5-2.5 μm (for B.1.351).

### Image processing and 3D reconstructions

Drift correction for cryo-EM images was performed using MotionCor2 (*44*), and contrast transfer function (CTF) was estimated by CTFFIND4 (*45*) using motion-corrected sums without dose-weighting. Motion corrected sums with dose-weighting were used for all other image processing. RELION3.0.8 and crYOLO (*46*) were used for particle picking, 2D classification, 3D classification and refinement procedure. For the B.1.1.7 sample, approximately 3,000 particles were manually picked for each protein sample and subjected to 2D classification to generate the templates for automatic particle picking. After manual inspection of auto-picked particles, a total of 2,325,106 particles were extracted from 27,950 images. The selected particles were subjected to 2D classification, giving a total of 877,530 good particles. A low-resolution negative-stain reconstruction of the D614 S trimer (*22*) was low-pass filtered to 30Å resolution and used as an initial model for 3D classification with C1 symmetry. Three major classes showed clear structural features were subjected to another round of 3D classification with C1 symmetry, giving another three major classes. These three classes were joined together and re-extract to do one round of 3D auto-refinement, following by CTF Refinement, Particle Polishing, and another round of 3D auto-refinement, giving a map with 271,387 particles at 3.3Å resolution. Third round of signal-subtraction 3D classification without alignment at the apex region of the S trimer were performed to further classify, produced five different major classes, representing the closed, three RBD-down conformation, three of the one RBD-up conformation and two RBD-up conformation, respectively. The five classes containing 13,919, 77,942, 7,368, 119,338, 41,138 particles, respectively, were then subjected to another round of 3D auto-refinement with C1 (one RBD-up and two RBD-up) and C3 (closed) symmetry using an overall mask, resulting in five final reconstructions at 4.0Å, 3.3Å, 4.3Å, 3.2Å and 3.1Å resolutions, respectively. Several rounds of 3D autorefinement was done for each then by adding different size of mask at the apex region to further improve the local resolution. The best map from each class was used for model building.

For the B.1.351 sample, after manual inspection of auto-picked particles, a total of 4,875,287 particles were extracted from 33,931 images. The selected particles were subjected to 2D classification, giving a total of 1,380,218 good particles. The low-resolution negative-stain reconstruction of the sample was low-pass-filtered to 40Å as an initial model for 3D classification with C1 symmetry. Two major classes with 224,988 and 301,967 particles that showed clear structural features were subjected to the second round of 3D classification with C1 symmetry. One class showing a 3-RBD down conformation with 124,042 particles was selected for auto-Refinement, giving a reconstruction at 4.2Å resolution.

Third round of signal-subtraction 3D classification without alignment at the apex region of the S trimer were performed to further classify, produced three different major classes, representing the closed, three RBD-down conformation, two of the one RBD-up conformation, respectively. The closed state classes containing 53,772 particles were then subjected to another round of 3D auto-refinement with C1 symmetry using an overall mask, resulting in final reconstruction at 4.5Å resolution. Several rounds of 3D auto-refinement were performed by adding a different size of mask at the apex region to further improve the local resolution. The best map from each class was used for model building.

For the RBD-up class of the B.1.351 sample, data processing was primarily carried out in cryoSPARC (*37*). Initially picked particles were subjected to sequential 2D classifications within RELION and cryoSPARC on downsampled data (3.3Å/pix) to give 994,071 particles. Two rounds of heterogeneous 3D refinement in cryoSPARC resulted in 552,519 particles being selected. Following downsampling to 1.65Å/pix, subsequent 3D heterogeneous refinement resulted in 503,181 particles corresponding to the open conformation. After un-binning to 0.825Å/pix these particles underwent Non-Uniform refinement, including cycles of CTF parameter refinement, resulting in a final resolution of 3.13Å. Particle polishing in RELION, and further Non-Uniform refinement including cycles of CTF parameter refinement, resulted in a final resolution of 2.95Å with no additional new features, compared to the unpolished map.

Reported resolutions are based on the gold-standard Fourier shell correlation (FSC) using the 0.143 criterion. All density maps from RELION were corrected from the modulation transfer function of the K3 detector and then sharpened by applying a temperature factor that was estimated using post-processing in the program. Local resolution was determined using RELION with half-reconstructions as input maps.

### Model building

The initial templates for model building used the stabilized SARS-CoV-2 S ectodomain trimer structure (PDB ID 7KRQ and PDB ID 7KRR) for both the B.1.351 and B.1.1.7 variant spike protein prefusion conformation. Several rounds of manual building were performed in Coot (*47*). The maps with a resolution lower than 4.0Å were primarily modeled manually in coot and by rigid body fitting, as the local resolution of many regions is higher than 4.0Å. The model was then refined in Phenix (*48*) against the 3.1Å (closed), 3.2Å, 3.3Å, 4.0Å (one RBD-up) and 4.3Å (two RBD-up) cryo-EM maps of the B.1.1.7 variant, and refined in Phenix against the 3.1Å (open), 4.5Å (closed) cryo-EM maps of the B.1.351 variant. Iteratively, refinement was performed in both Phenix (real space refinement) and ISOLDE (*49*), and the Phenix refinement strategy included minimization_global, local_grid_search, and adp, with rotamer, Ramachandran, and reference-model restraints, using 7KRQ and 7KRR as the reference model. The refinement statistics are summarized in Table S2. Structural biology applications used in this project were compiled and configured by SBGrid (*50*).

