## Supplemental Figures and Tables for "Structural basis for enhanced infectivity and immune evasion of SARS-CoV-2 variants"

#### Supplementary materials

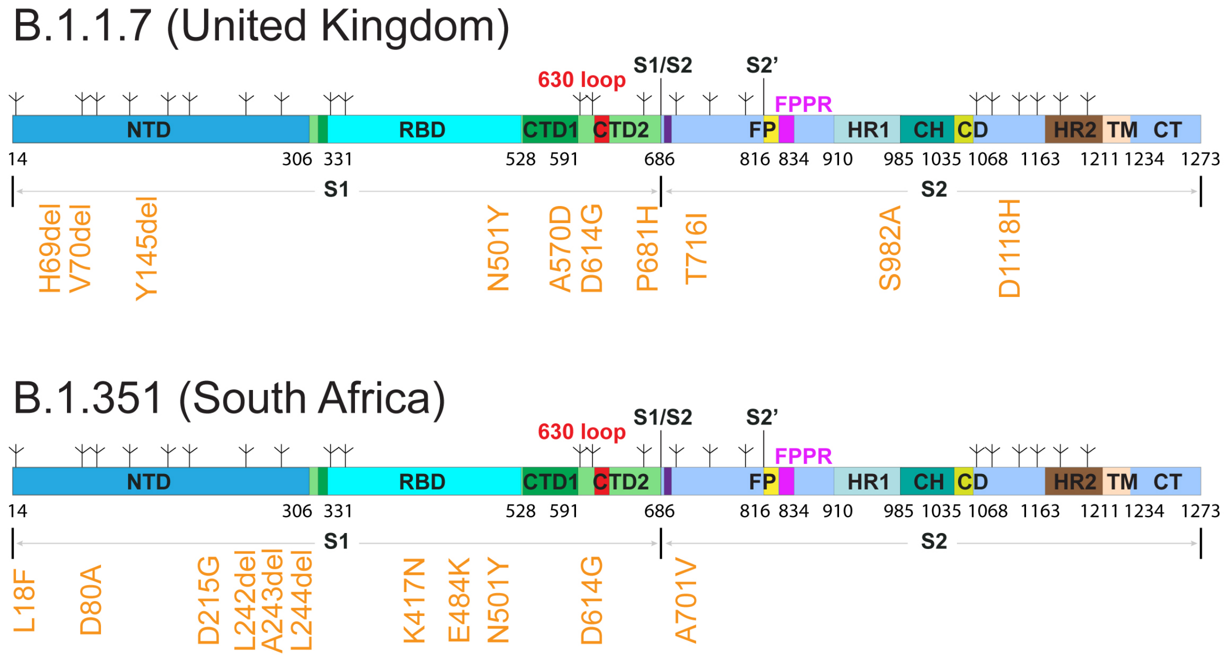


**Figure S1. Schematic representation of the full-length SARS-CoV-2 spike (S) from the B.1.1.7 and B.1.351 variants.** Segments of S1 and S2 include: NTD, N-terminal domain; RBD, receptor-binding domain; CTD1, C-terminal domain 1; CTD2, C-terminal domain 2; 630 loop; S1/S2, S1/S2 cleavage site; S2', S2' cleavage site; FP, fusion peptide; FPPR, fusion peptide proximal region; HR1, heptad repeat 1; CH, central helix region; CD, connector domain; HR2, heptad repeat 2; TM, transmembrane anchor; CT, cytoplasmic tail; and tree-like symbols for glycans. Positions of all mutations (from the amino-acid sequence of Wuhan-Hu-1) are shown in orange text.

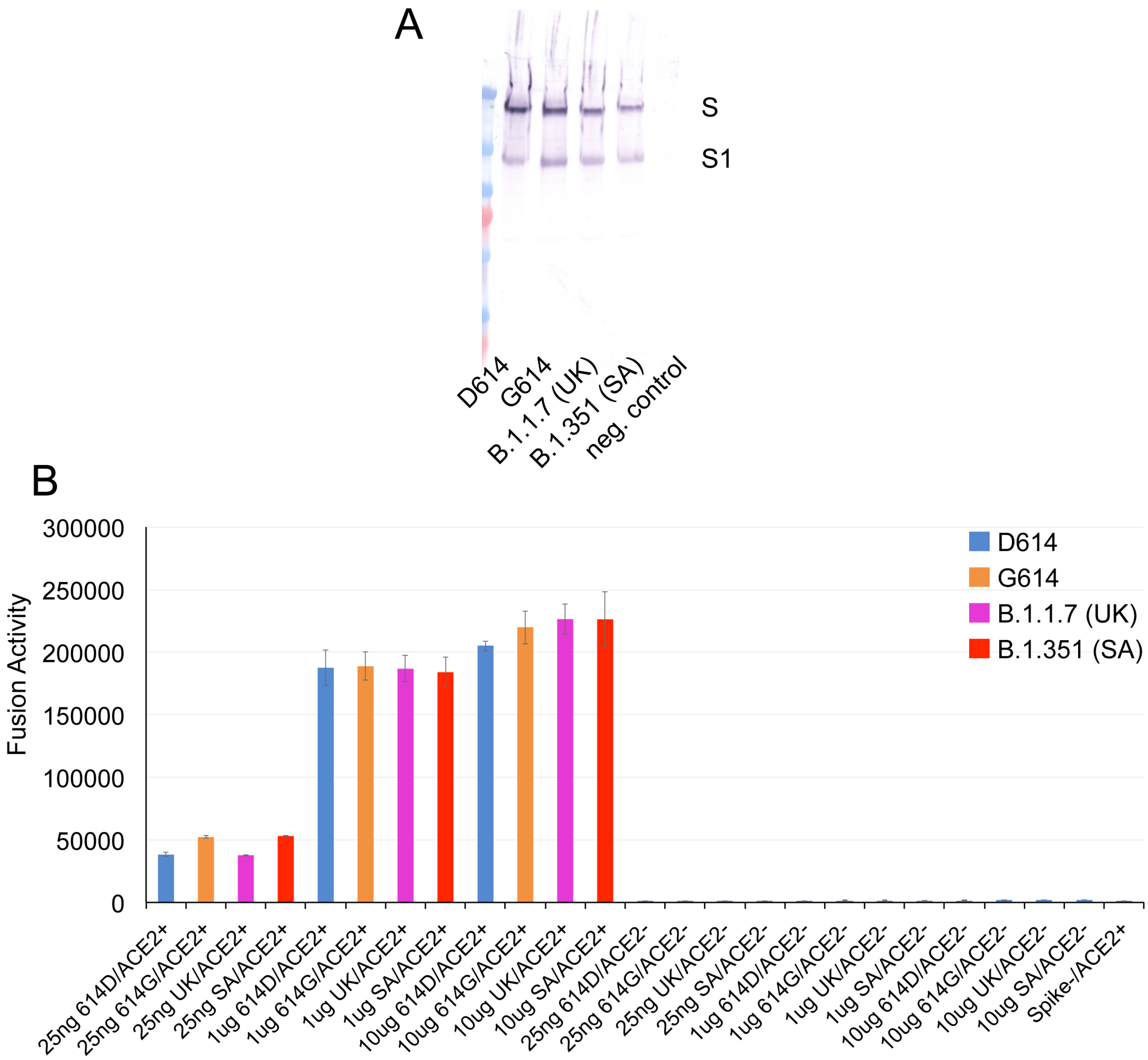


**Figure S2. Expression and cell-cell fusion of SARS-CoV-2 variants.** (**A**) Expression and processing of the full-length S constructs in HEK293 cells. S samples prepared from HEK293 cells transiently transfected with 10 μg of the full-length S expression plasmids were detected by anti-RBD polyclonal antibodies. Bands for the uncleaved S and S1 fragment are indicated. (**B**) HEK293T cells transfected with either the untagged full length S protein expression plasmids were fused with ACE2-expressing cells. Cell-cell fusion led to reconstitution of α and ω fragments of β-galactosidase yielding an active enzyme, and the fusion activity was then quantified by a chemiluminescent assay. No ACE2 and no S were negative controls.

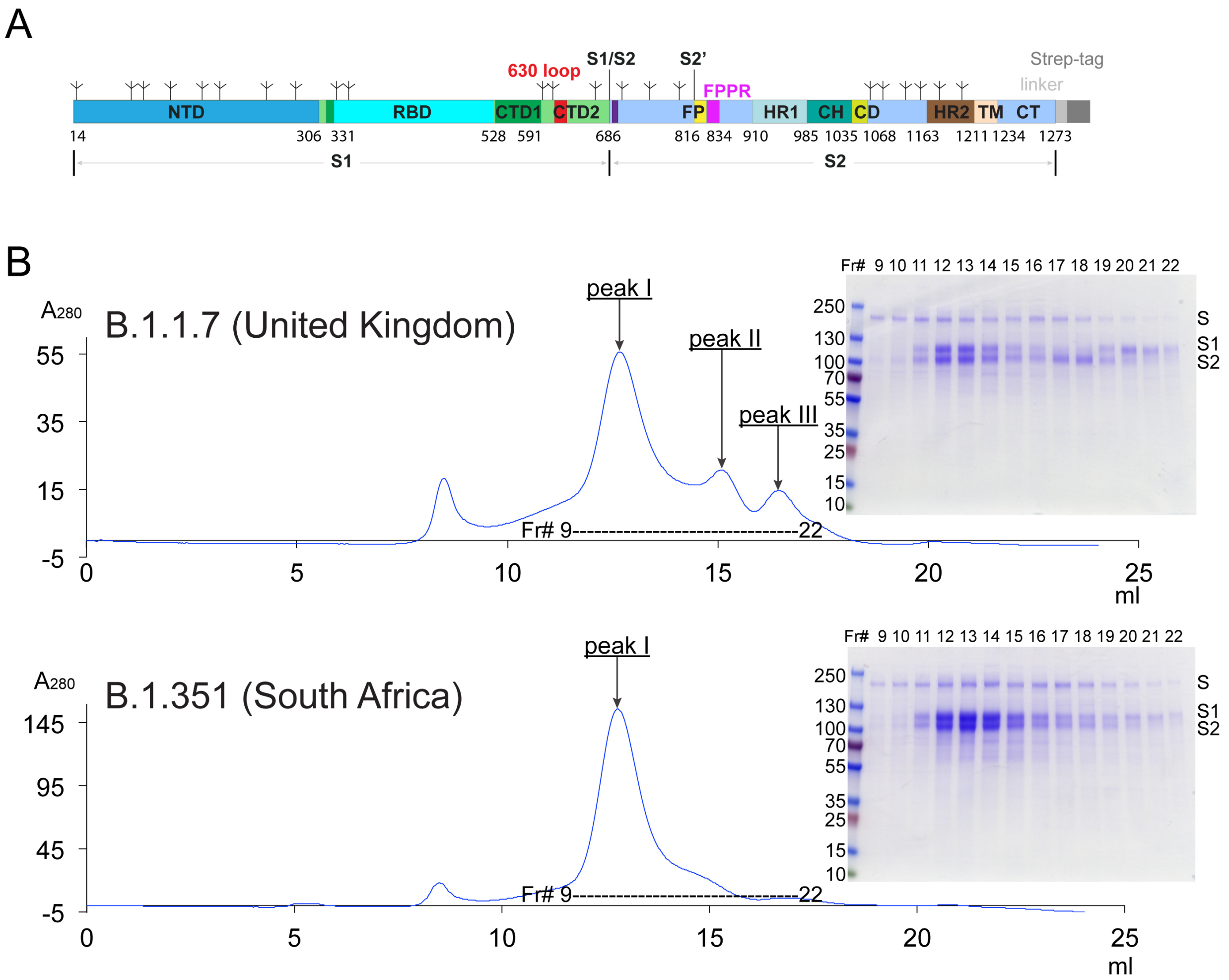


**Figure S3. Production of full-length S protein from the B.1.1.7 and B.1.351 variants.** (**A**) A strep-tag was fused to the C-terminus of the full-length S protein by a flexible linker. (**B**) The full-length S proteins were extracted and purified in detergent DDM, and further resolved by gel-filtration chromatography on a Superose 6 column. Peak I, the prefusion S trimer; peak II, the postfusion S2 trimer; and peak III, the dissociated monomeric S1. Inset, peak fractions were analyzed by Coomassie stained SDS-PAGE. Labeled bands are S, S1 and S2. Fr#, fraction number.

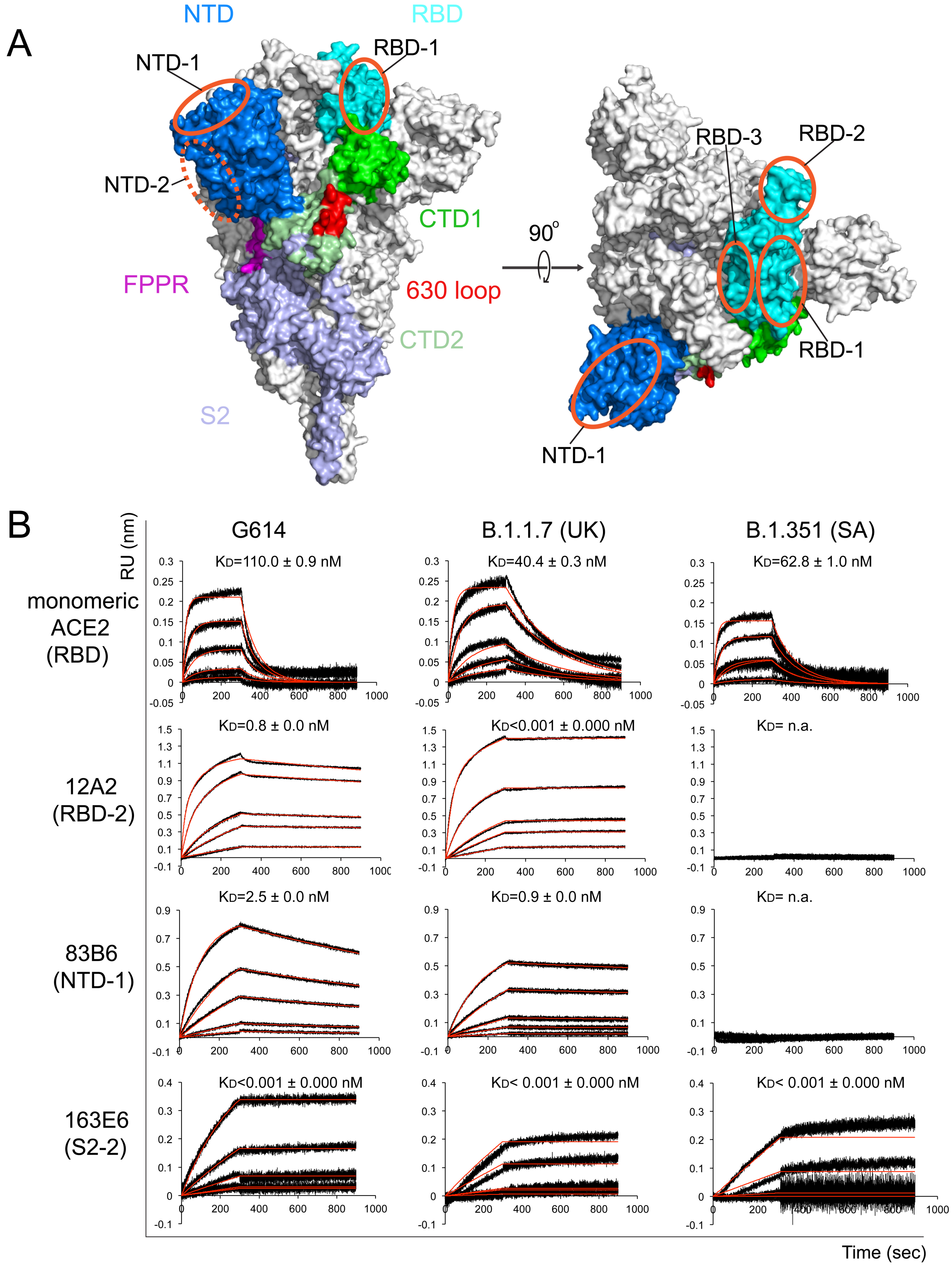


**Figure S4. Additional antigenic properties of the purified full-length SARS-CoV-2 S proteins**. (**A**) Antibody competition clusters as described in ref(*35*). Surface regions of the S trimer targeted by antibodies on S1 are highlighted by orange ellipses, including RBD-1, RBD-2, RBD-3, NTD-1 and NTD-2. The exact location of NTD-2 is uncertain and therefore marked with a dashed line. (**B**) Binding analysis of the prefusion S trimers from G614, B.1.1.7 and B.1.351 variants with soluble ACE2 constructs was performed by bio-layer interferometry (BLI). For ACE2 binding, the purified S proteins were immobilized on AR2G biosensors and dipped into the wells containing ACE2 at various concentrations. For antibody binding, various antibodies were immobilized to AHC biosensors and dipped into the wells containing each purified S protein at different concentration. Binding kinetics were evaluated using a 1:1 Langmuir model except for antibody 12A2 targeting the RBD-2, which was analyzed by a bivalent binding model. The sensorgrams are in black and the fits in red. RU, response unit. Binding constants are also summarized here and in Table S1. All experiments were repeated at least twice with essentially identical results.

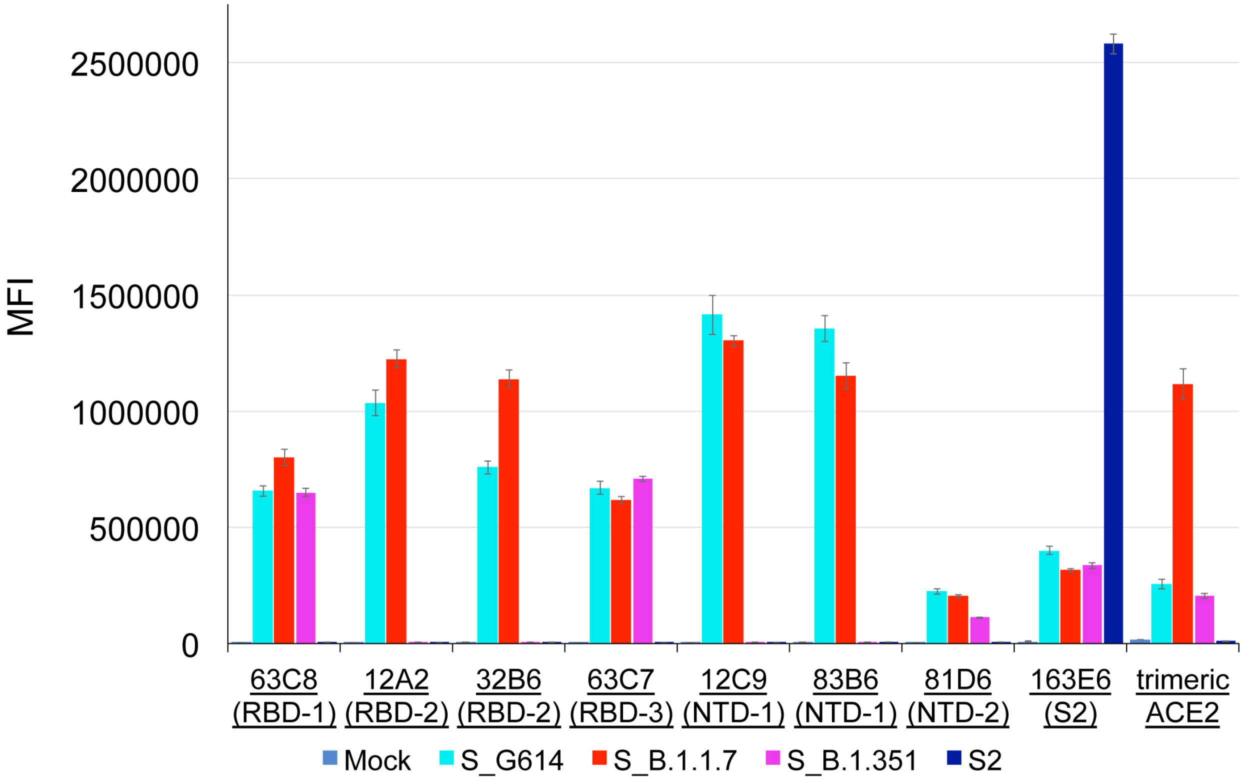


**Figure S5. Antigenic properties of the cell-surface S proteins assessed by flow cytometry.** Antibody and ACE2 binding to the full-length S proteins of the G614, B.1.1.7 and B.1.351 variants, as well as an S2 construct expressed on the cell surfaces analyzed by flow cytometry. The antibodies and their targets are indicated. A designed ACE2-based inhibitor $ACE2_{615}$-foldon-T27W was used for detecting receptor binding (*41*). MFI, mean fluorescent intensity. The flow cytometry assays were repeated three times with essentially identical results.

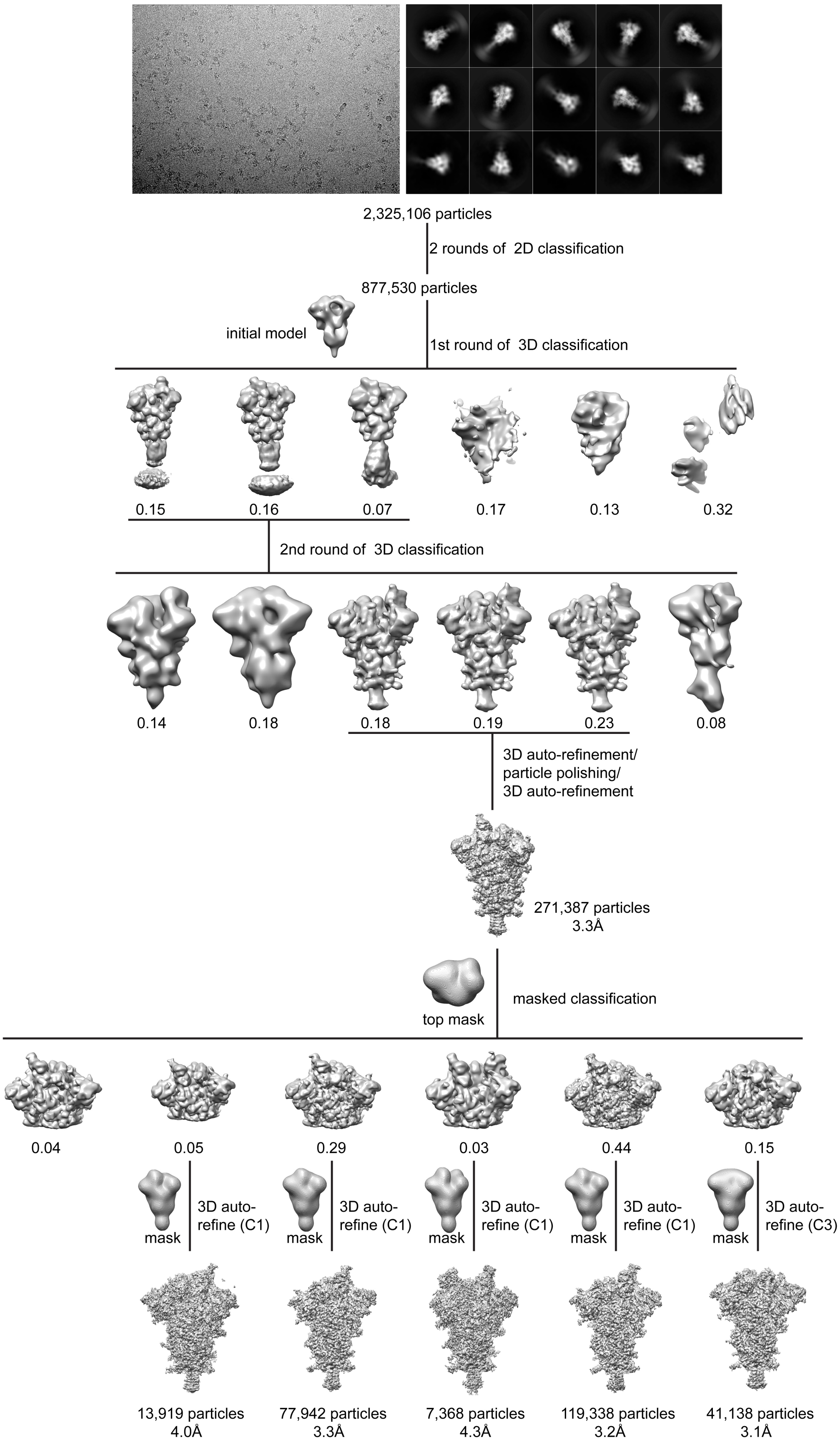

2,325,106 particles
2 rounds of 2D classification
877,530 particles
initial model
1st round of 3D classification
0.15
0.16
0.07
0.17
0.13
0.32
2nd round of 3D classification
0.14
0.18
0.18
0.19
0.23
0.08
3D auto-refinement/
particle polishing/
3D auto-refinement
271,387 particles
3.3Å
top mask
masked classification
0.04
0.05
0.29
0.03
0.44
0.15
mask
3D auto-refine (C1)
mask
3D auto-refine (C1)
mask
3D auto-refine (C1)
mask
3D auto-refine (C1)
mask
3D auto-refine (C3)
13,919 particles
4.0Å
77,942 particles
3.3Å
7,368 particles
4.3Å
119,338 particles
3.2Å
41,138 particles
3.1Å

**Figure S6. Cryo-EM analysis of the B.1.1.7 S trimer.** Top, representative micrograph**,** and 2D averages (box dimension: 396Å) of the cryo-EM particle images of the B.1.1.7 S trimer. Bottom, data processing workflow for structure determination.

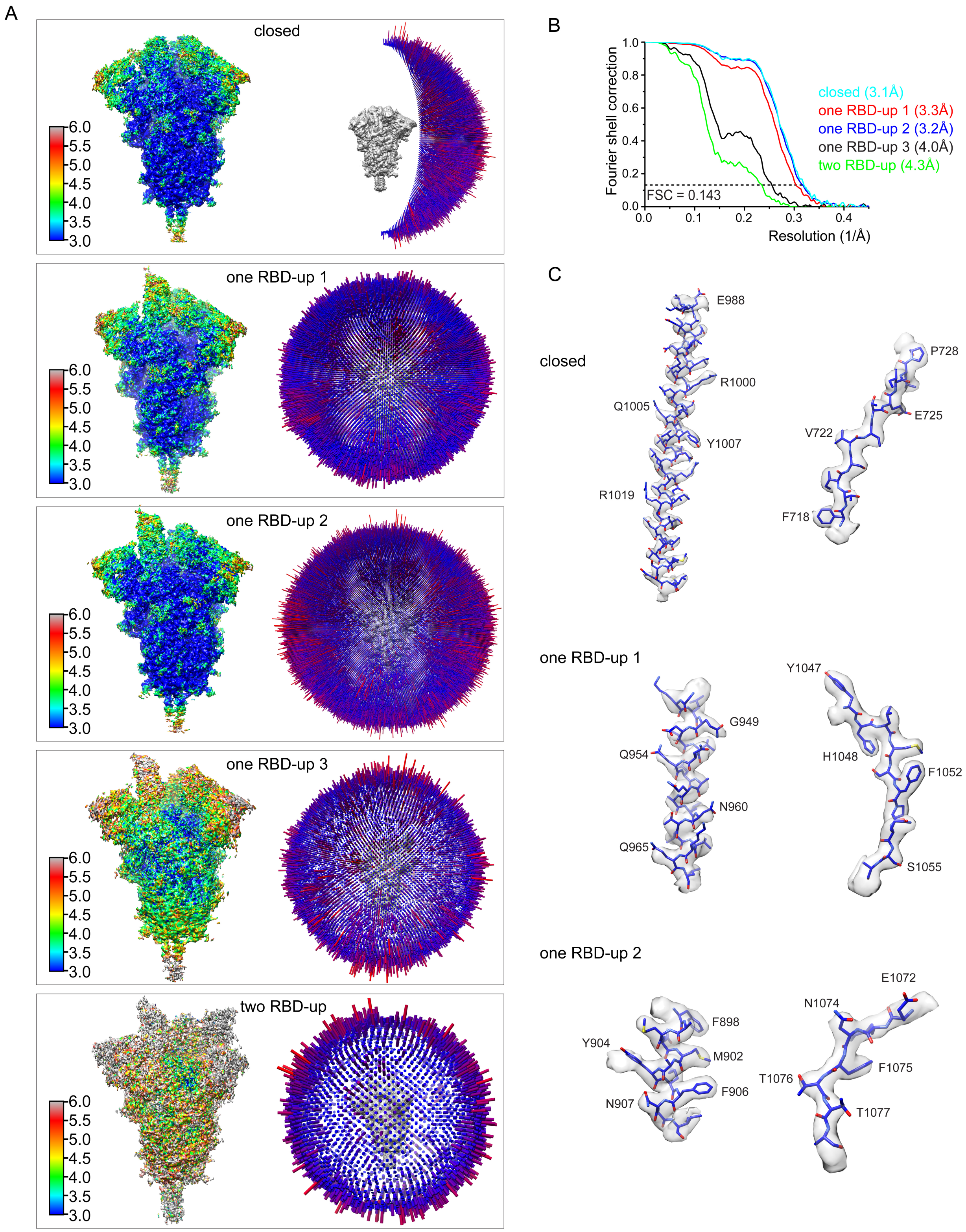


**Figure S7. Analysis of the B.1.1.7 S trimer structure.** (A) 3D reconstructions of the B.1.1.7 S trimer preparation in the closed, three one RBD-up and a two RBD-up conformations, respectively, are colored according to local resolution estimated by RELION. Angular distribution of the cryo-EM particles used in each reconstruction is shown in the side view of the EM map. (B) Gold standard FSC curves of the three refined 3D reconstructions of the B.1.1.7 S trimer. (C) Representative density in gray surface from EM maps with a resolution better than 3.5Å.

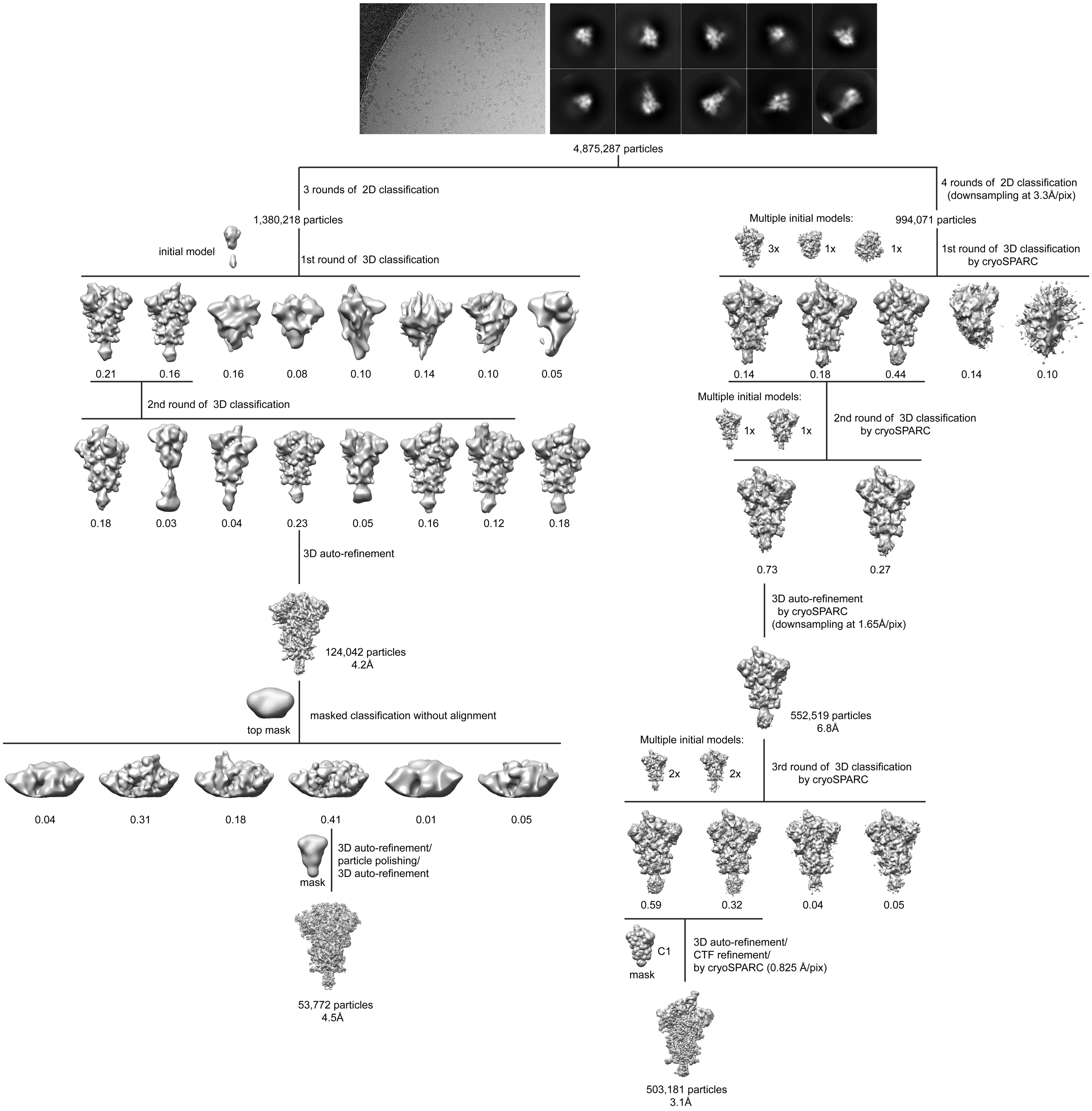


**Figure S8. Cryo-EM analysis of the B.1.351 S trimer.** Top, representative micrograph**,** and 2D averages (box dimension: 396Å) of the cryo-EM particle images of the B.1.351 S trimer. Bottom, data processing workflow for structure determination.

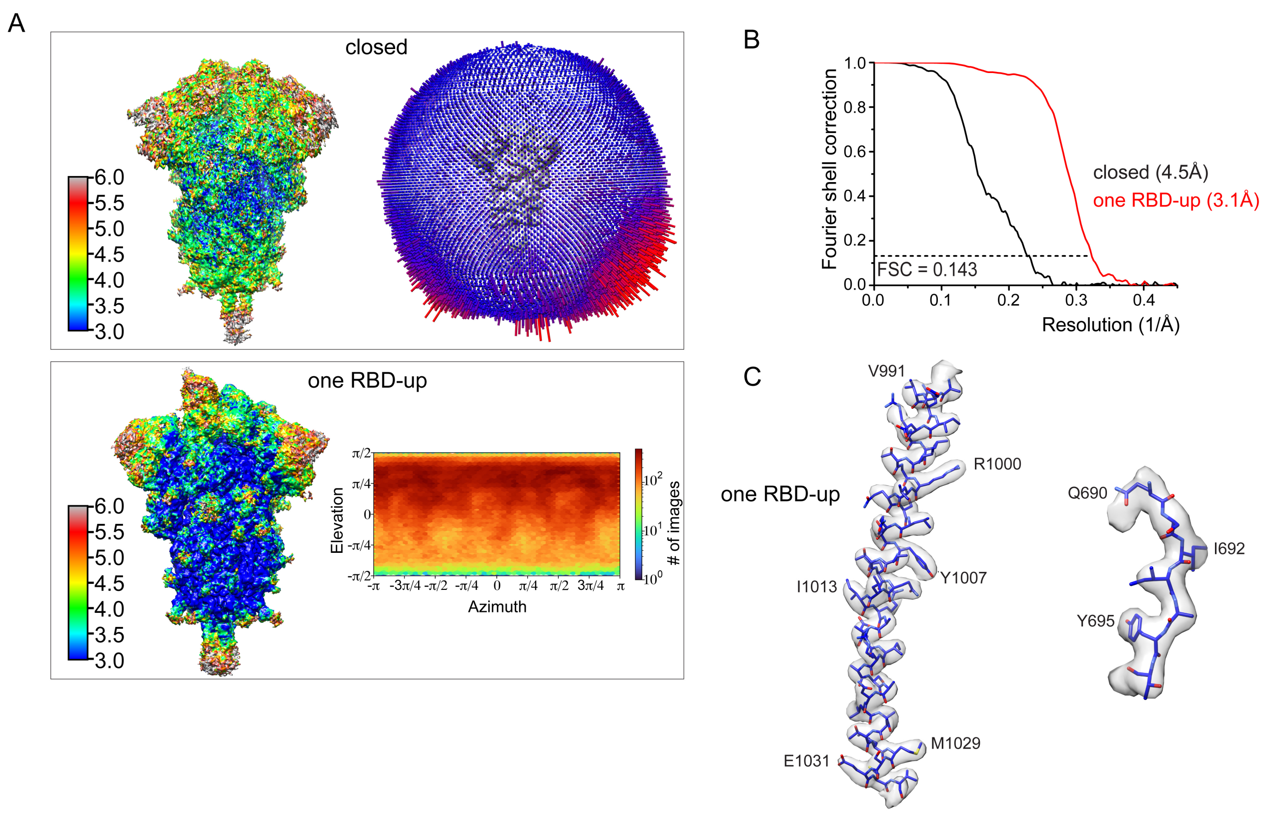


**Figure S9. Analysis of the B.1.351 S trimer structure.** (A) 3D reconstructions of the B.1.351 S trimer preparation in the closed and one RBD-up conformations, respectively, are colored according to local resolution estimated by RELION. Angular distribution of the cryo-EM particles used in reconstruction for the closed conformation is shown in the side view of the EM map. The RBD-up conformation was reconstructed using cryoSPARC and the output from the program for particle distribution is shown. (B) Gold standard FSC curves of the refined 3D reconstructions of the B.1.351 S trimer. (C) Representative density in gray surface from EM map of the one RBD-up structure.

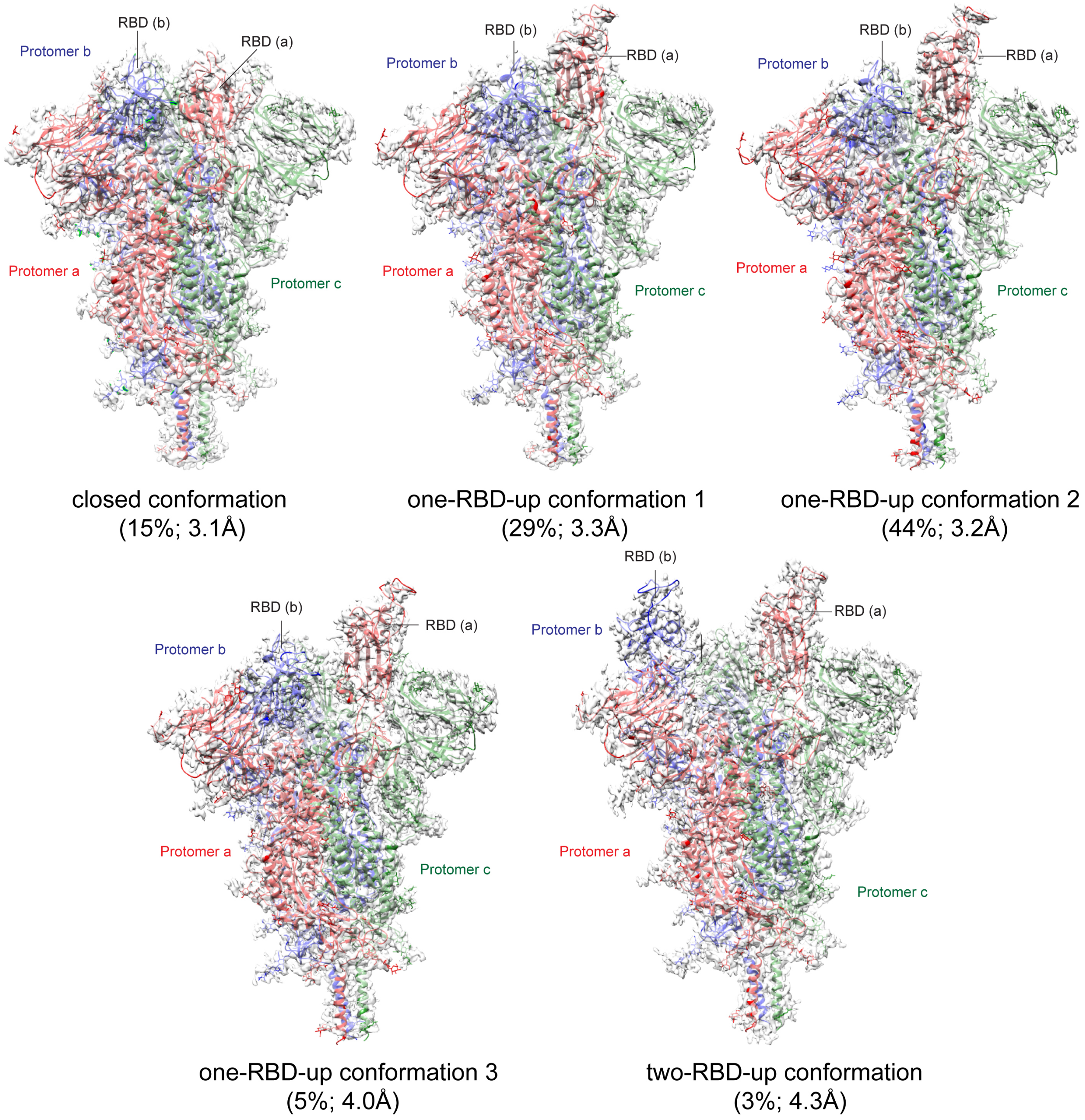


**Figure S10. Cryo-EM structures of the full-length S protein of the B.1.1.7 variant**. Five structures of the B.1.1.7 S trimer, representing a closed prefusion conformation, three distinct RBD-up conformations, and a two RBD-up conformation, were modeled based on corresponding cryo-EM density maps at 3.1-4.3Å resolution. The maps with a resolution lower than 4.0Å were primarily modeled manually in coot and by rigid body fitting, as the local resolution of many regions is higher than 4.0Å. Three protomers (a, b, c) are colored in red, blue and green, respectively. RBD locations are indicated. Particle percentage for each class in the data processing is also indicated, but it may not accurately reflect the conformation distribution of the S trimer in solution.

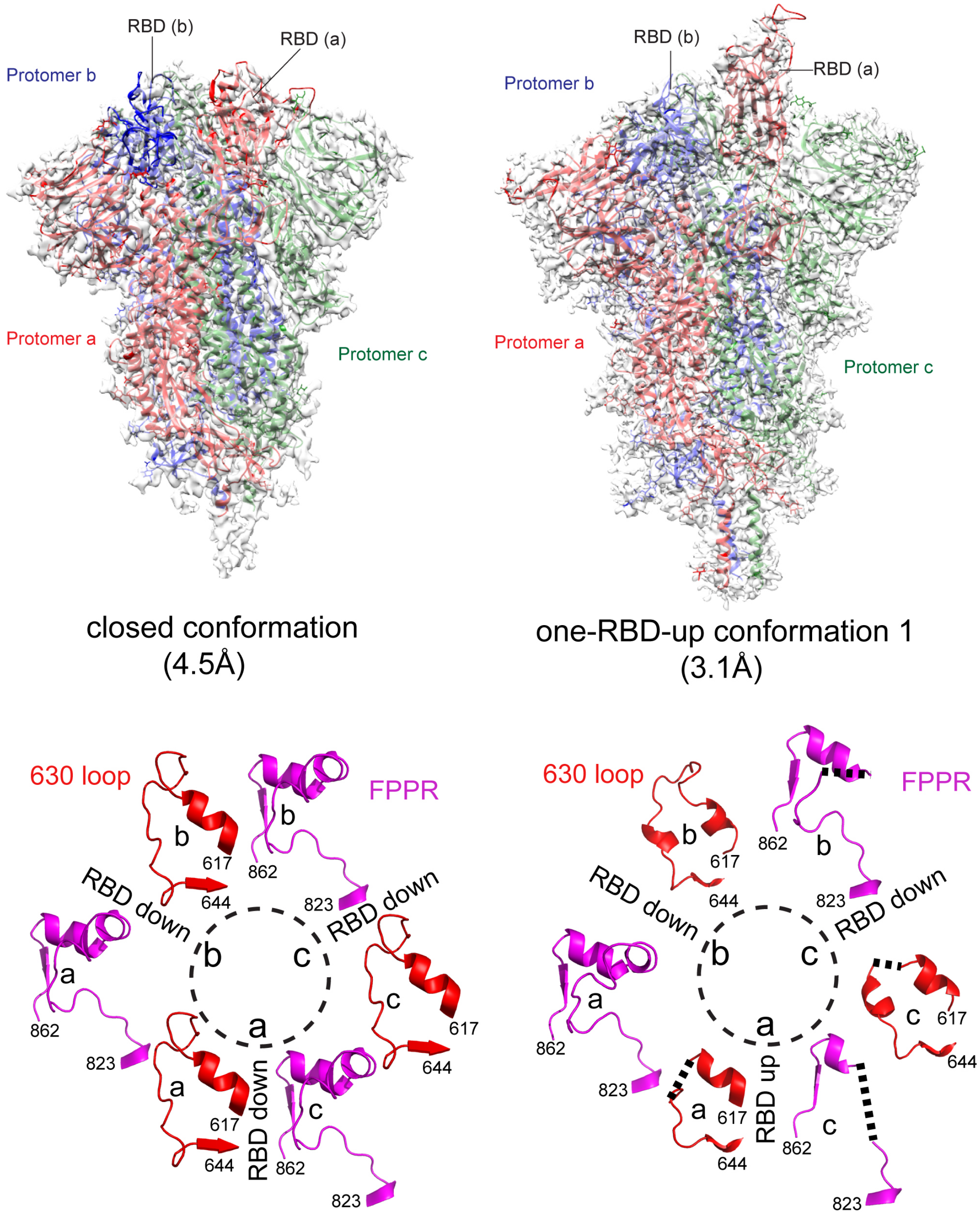


**Figure S11. Cryo-EM structures of the full-length S protein of the B.1.351 variant**. Top, two structures of the B.1.351 S trimer, representing a closed prefusion conformation, and an RBD-up conformation were modeled based on corresponding cryo-EM density maps at 4.5Å and 3.1Å resolution, respectively. The map of the closed conformation was modeled manually in coot and also by rigid body fitting, as the local resolution of many regions is higher than 4.5Å. Three protomers (a, b, c) are colored in red, blue and green, respectively. RBD locations are indicated. Bottom, structures of three segments (residues 617-644) containing the 630 loop in red and three segments (residues 823-862) containing the FPPR in magenta from all three protomers (a, b and c) are shown for the B.1.1.7 trimer. Position of each RBD is indicated. Dashed lines indicate gaps.

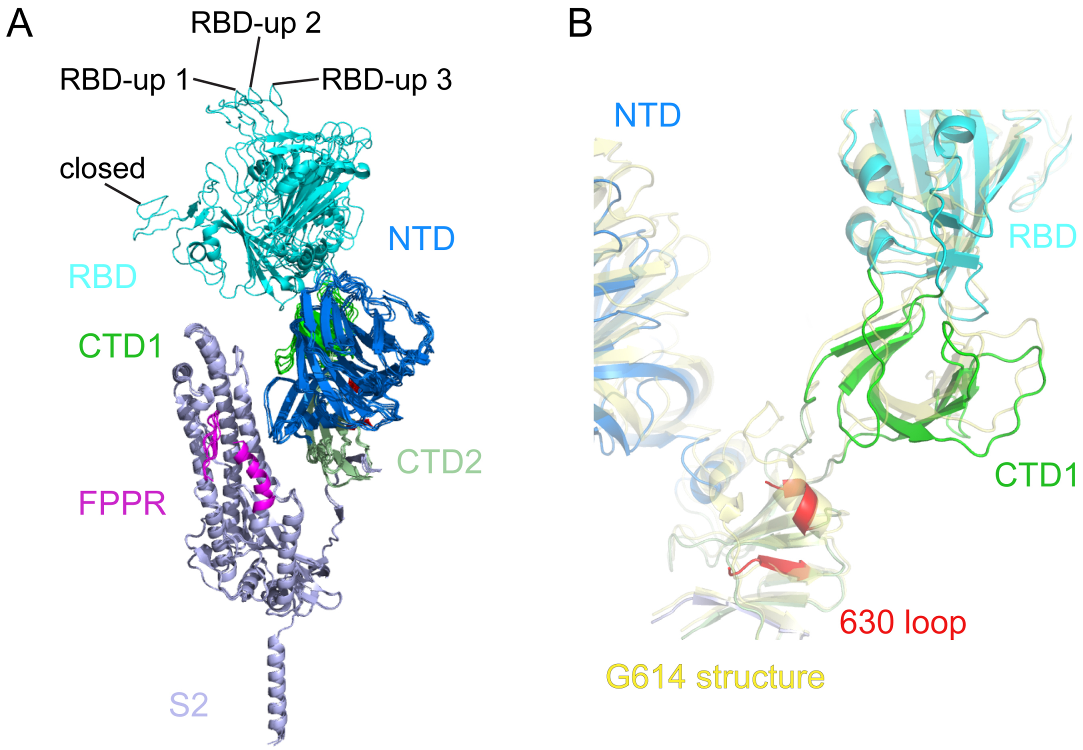


**Figure S12. Superposition of the B.1.1.7 trimer structures and the G614 structure.** (**A**) Side views of superposition of the closed conformation and three distinct one RBD-up conformations of the B.1.1.7 S in ribbon, align by the S2 portion. The positions of the RBD-down and three different RBD-up conformations are indicated. (**B**) Comparison of the CTD1 in the B.1.1.7 (various colors) and G614 trimer (yellow) structures, when aligned by S2.

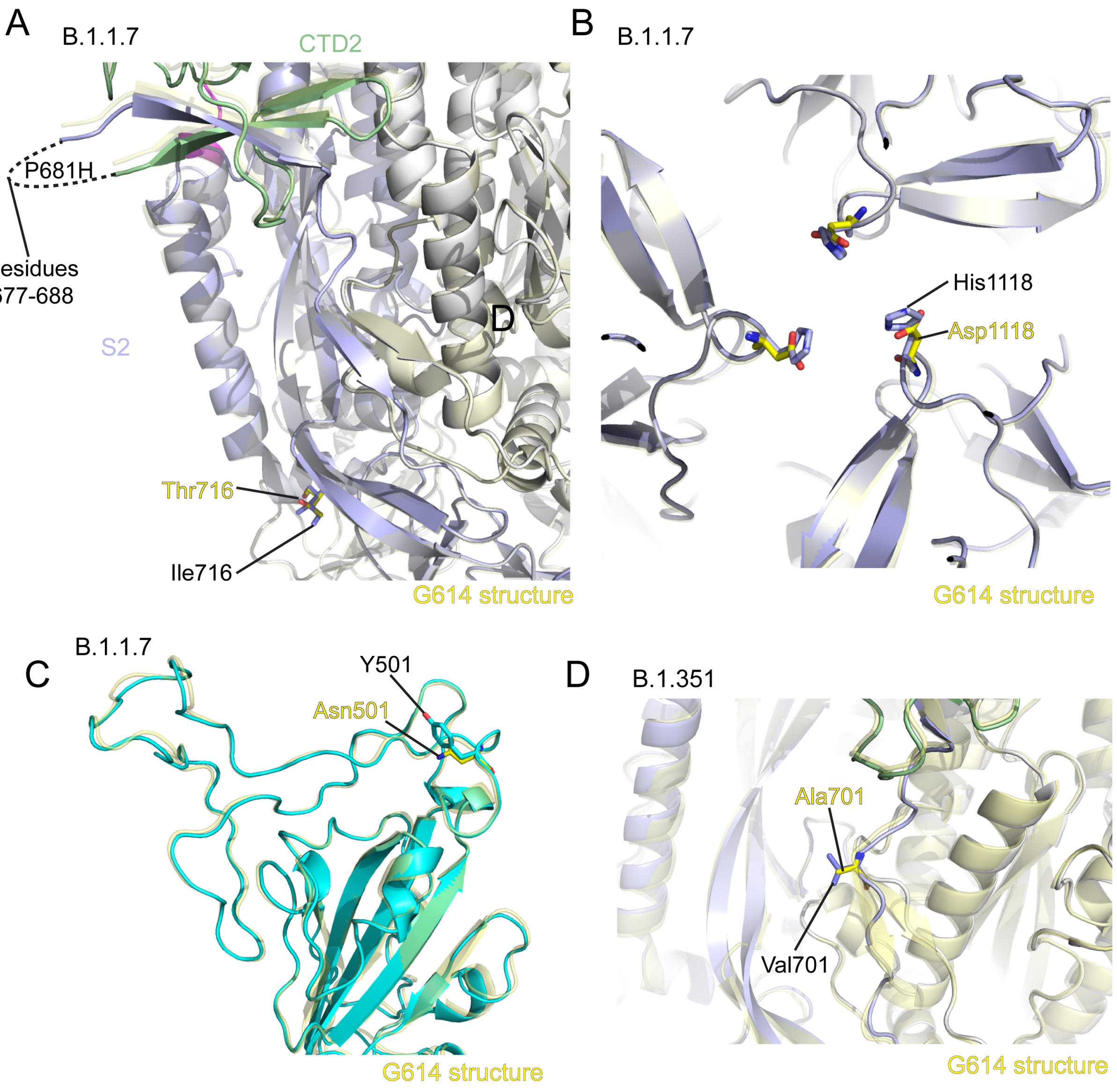


**Figure S13. Structural impact of the mutations in the variants**. (**A-C**) Views of superposition of the structure of the B.1.1.7 S trimer in ribbon representation with the structure of the G614 S in yellow, showing the regions near mutations N501Y, P681H, T716I and D1118H. (**D**) A view of superposition of the structure of the B.1.351 S trimer in ribbon representation with the structure of the G614 S in yellow, showing the region near the mutation A701V. All mutations are indicated and highlighted as sticks.

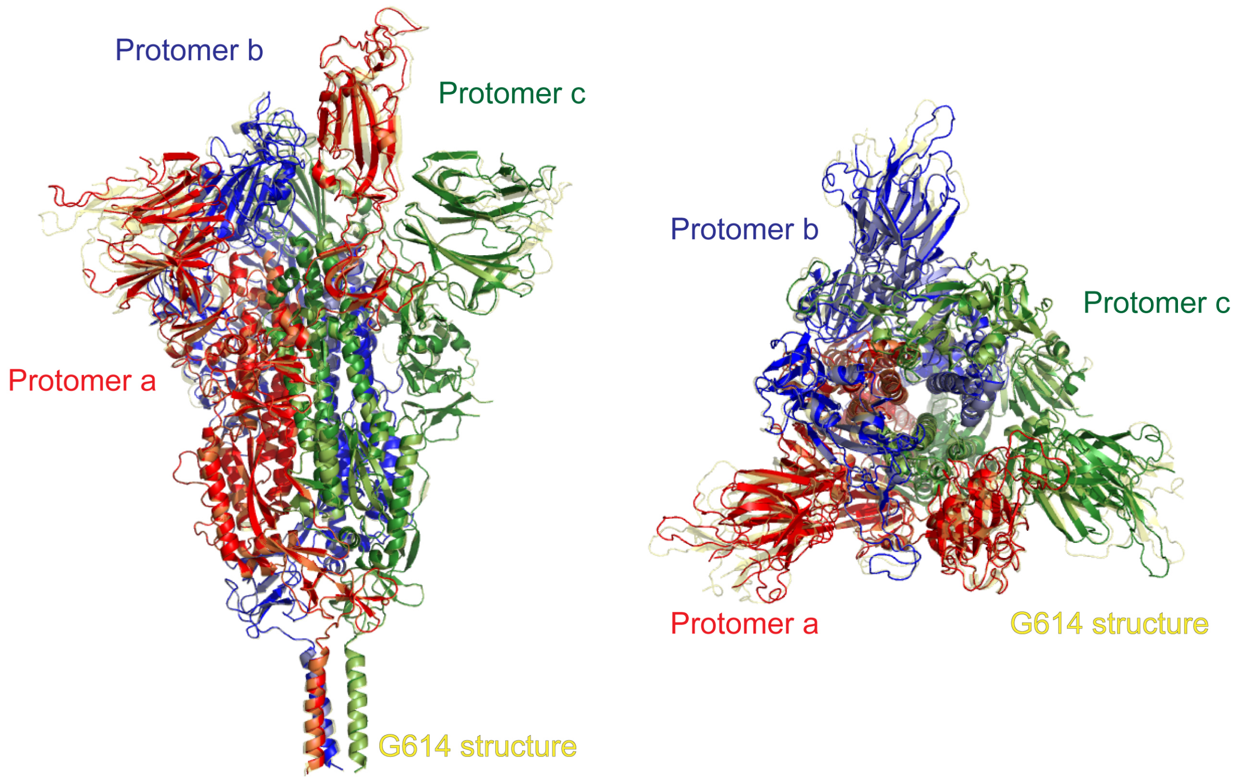


**Figure S14. Superposition of the structures of the B.1.351 and G614 S trimers in the one RBD-up conformation.** The structures of the B.1.351 and G614 S (PDB ID: 7KRR) trimers in the one RBD-up conformation are aligned by the invariant S2. Three protomers (a, b, c) are colored in red, blue and green, respectively.

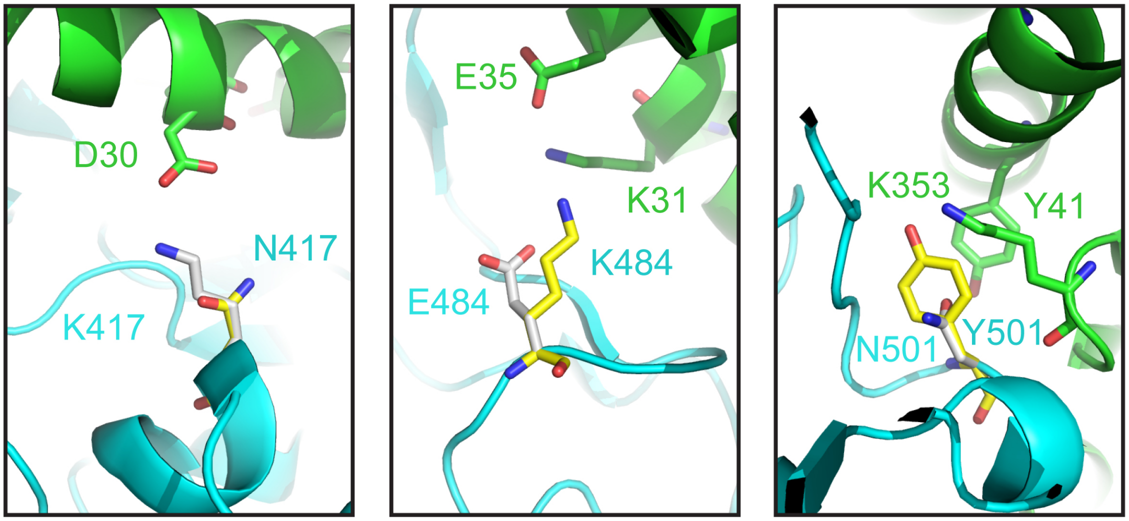


**Figure S15. Modeled interface between the RBD and ACE2.** The interface between ACE2 in ribbon diagram in green and RBD in cyan from the complex structure (PDB ID: 6M0J; ref(*17*)). Modeled K417N, E484K and N501Y are shown as sticks.

### Table S1. Binding constants of S-ACE2 interaction

| | | $K_D$ (M) | $K_D$ Error | ka (1/Ms) | ka2 | ka Error | ka2 Error | kdis (1/s) | kdis2 | kdis Error | kdis2 Error |
|---|---|---|---|---|---|---|---|---|---|---|---|
| **ACE2-Fc (RBD)** | G614 | 1.09E-08 | 6.41E-10 | 4.91E+04 | 5.89E+00 | 2.08E+03 | 2.63E+02 | 5.34E-04 | 9.76E-01 | 2.19E-05 | 4.34E+01 |
| | B.1.1.7(UK) | 2.35E-09 | 1.78E-10 | 8.23E+04 | 3.49E-01 | 3.97E+03 | 3.15E-01 | 1.94E-04 | 2.94E-02 | 1.13E-05 | 2.77E-02 |
| | B.1.351(SA) | 1.33E-08 | 7.25E-10 | 5.62E+04 | 1.08E+01 | 1.22E+03 | 1.40E+02 | 7.47E-04 | 3.98E-01 | 3.74E-05 | 5.12E+00 |
| **Monomeric ACE2 (RBD)** | G614 | 1.10E-07 | 9.20E-10 | 1.10E+05 | | 7.73E+02 | | 1.20E-02 | | 5.43E-05 | |
| | B.1.1.7(UK) | 4.04E-08 | 3.51E-10 | 8.65E+04 | | 7.19E+02 | | 3.49E-03 | | 8.84E-06 | |
| | B.1.351(SA) | 6.28E-08 | 9.74E-10 | 1.31E+05 | | 1.95E+03 | | 8.22E-03 | | 3.58E-05 | |
| **63C8 (RBD-1)** | G614 | 1.03E-08 | 5.38E-11 | 9.10E+04 | | 4.40E+02 | | 9.41E-04 | | 1.81E-06 | |
| | B.1.1.7(UK) | 1.72E-09 | 8.60E-12 | 1.63E+05 | | 4.27E+02 | | 2.81E-04 | | 1.19E-06 | |
| | B.1.351(SA) | 1.74E-08 | 2.22E-10 | 5.58E+04 | | 6.88E+02 | | 9.70E-04 | | 2.13E-06 | |
| **32B6 (RBD-2)** | G614 | 9.38E-10 | 8.85E-12 | 2.97E+05 | 2.42E-01 | 1.58E+03 | 2.82E-02 | 2.79E-04 | 4.36E-02 | 2.17E-06 | 4.94E-03 |
| | B.1.1.7(UK) | 1.09E-10 | 1.45E-11 | 1.64E+05 | 1.02E-01 | 2.04E+03 | 1.78E-02 | 1.79E-05 | 5.37E-03 | 2.37E-06 | 3.75E-04 |
| | B.1.351(SA) | N.D. | N.D. | N.D. | N.D. | N.D. | N.D. | N.D. | N.D. | N.D. | N.D. |
| **12A2 (RBD-2)** | G614 | 7.53E-10 | 1.10E-11 | 3.03E+05 | 3.63E-01 | 2.58E+03 | 8.78E-02 | 2.28E-04 | 5.66E-02 | 2.72E-06 | 1.32E-02 |
| | B.1.1.7(UK) | <1.0E-12 | 2.26E-11 | 1.40E+05 | 1.09E-01 | 2.13E+03 | 3.28E-02 | <1.0E-07 | 4.93E-03 | 3.16E-06 | 7.14E-04 |
| | B.1.351(SA) | N.D. | N.D. | N.D. | N.D. | N.D. | N.D. | N.D. | N.D. | N.D. | N.D. |
| **63C7 (RBD-3)** | G614 | 1.13E-08 | 4.21E-11 | 1.81E+05 | | 6.39E+02 | | 2.05E-03 | | 2.46E-06 | |
| | B.1.1.7(UK) | 1.66E-09 | 1.33E-11 | 1.07E+05 | | 3.94E+02 | | 1.79E-04 | | 1.27E-06 | |
| | B.1.351(SA) | 1.19E-08 | 3.20E-11 | 1.75E+05 | | 4.48E+02 | | 2.09E-03 | | 1.76E-06 | |
| **12C9 (NTD-1)** | G614 | 7.31E-09 | 3.64E-11 | 7.02E+04 | | 3.08E+02 | | 5.13E-04 | | 1.22E-06 | |
| | B.1.1.7(UK) | 3.89E-10 | 2.95E-12 | 3.25E+05 | | 6.22E+02 | | 1.26E-04 | | 9.26E-07 | |
| | B.1.351(SA) | N.D. | N.D. | N.D. | | N.D. | | N.D. | | N.D. | |
| **83B6 (NTD-1)** | G614 | 2.48E-09 | 7.39E-12 | 1.89E+05 | | 3.87E+02 | | 4.70E-04 | | 1.02E-06 | |
| | B.1.1.7(UK) | 9.21E-10 | 1.17E-11 | 1.21E+05 | | 4.44E+02 | | 1.11E-04 | | 1.35E-06 | |
| | B.1.351(SA) | N.D. | N.D. | N.D. | | N.D. | | N.D. | | N.D. | |
| **81D6 (NTD-2)** | G614 | 2.14E-09 | 1.33E-11 | 1.91E+05 | | 7.57E+02 | | 4.08E-04 | | 1.96E-06 | |
| | B.1.1.7(UK) | 2.10E-09 | 3.02E-11 | 6.52E+04 | | 4.80E+02 | | 1.37E-04 | | 1.69E-06 | |
| | B.1.351(SA) | 3.25E-09 | 3.28E-11 | 7.12E+04 | | 4.77E+02 | | 2.31E-04 | | 1.75E-06 | |
| **163E6 (S2-2)** | G614 | <1.0E-12 | 2.14E-12 | 7.18E+04 | | 5.84E+02 | | <1.0E-07 | | 1.53E-07 | |
| | B.1.1.7(UK) | <1.0E-12 | 2.41E-11 | 1.66E+04 | | 1.46E+03 | | <1.0E-07 | | 3.99E-07 | |
| | B.1.351(SA) | <1.0E-12 | 3.49E-09 | 2.90E+03 | | 2.55E+03 | | <1.0E-07 | | 1.01E-05 | |

**Table S2. Neutralization of the SARS-CoV-2 variants**

| **Antibody/ACE2 construct** | **Neutralization titer (µg/ml)** | | | | | |
|---|---|---|---|---|---|---|
| | **G614** | | **B.1.1.7 (UK)** | | **B.1.351 (South Africa)** | |
| | $IC_{50}$ | $IC_{80}$ | $IC_{50}$ | $IC_{80}$ | $IC_{50}$ | $IC_{80}$ |
| 63C8 (RBD-1) | 3.815 | 31.072 | 3.148 | 37.899 | 7.570 | 31.210 |
| 12A2 (RBD-2) | 0.020 | 0.080 | 0.047 | 0.204 | >50 | >50 |
| 32B6 (RBD-2) | 0.019 | 0.074 | 0.022 | 0.069 | >50 | >50 |
| 63C7 (RBD-3) | 8.081 | 44.520 | 2.067 | 17.671 | 11.450 | 48.346 |
| 12C9 (NTD-1) | 0.023 | 4.045 | 0.015 | 4.278 | >50 | >50 |
| 83B6 (NTD-1) | 0.189 | >50 | 4.541 | >50 | >50 | >50 |
| 81D6 (NTD-2) | >50 | >50 | >50 | >50 | >50 | >50 |
| 163E6 (S2-2) | >50 | >50 | >50 | >50 | >50 | >50 |
| ACE2-T27W-Fd | 0.138 | 0.774 | 0.028 | 0.157 | 0.082 | 0.557 |
| Positive serum pool_2 (1/x) | 753 | 132 | 711 | 103 | 41 | <20 |
| Normal Human Serum (1/x) | <20 | <20 | <20 | <20 | <20 | <20 |

**Table S3. Cryo-EM statistics.**

**EM data collection and reconstruction statistics**

| Protein | Full-length B.1.351 S protein | | Full-length B.1.1.7 S protein | | | | |
|---|---|---|---|---|---|---|---|
| Detergent | DDM | | DDM | | | | |
| Microscope | Titan Krios | | Titan Krios | | | | |
| Voltage(kV) | 300 | | 300 | | | | |
| Detector | Gatan K3 | | Gatan K3 | | | | |
| Magnification(nominal) | 105,000 | | 105,000 | | | | |
| Energy filter slit width (eV) | 20 | | 20 | | | | |
| Calibrated pixel size (Å/pix) | 0.825 | | 0.825 | | | | |
| Exposure rate ($e^-$/pix/sec) | 20.468 | | 20.21 | | | | |
| Frames per exposure | 50 | | 51 | | | | |
| Total electron exposure ($e^-/Å^2$) | 51.1 | | 53.4 | | | | |
| Exposure per frame ($e^-/Å^2$) | 1.022 | | 1.048 | | | | |
| Defocus range (µm) | -1.5, -2.5 | | -1.4, -2.3 | | | | |
| Automation software | SerialEM | | SerialEM | | | | |
| # of Micrographs used | 33,931 | | 27,950 | | | | |
| Particles extracted | 4,875,287 | | 2,325,106 | | | | |
| Particles after 2D classification | 1,380,218 | | 877,530 | | | | |
| Class | Closed | 1RBD-up | Closed | 1 RBD-up 1 | 1 RBD-up 2 | 1 RBD-up 3 | 2 RBD-up |
| Total # of refined particles | 53,772 | 503,181 | 41,138 | 77,942 | 119,338 | 13,919 | 7,368 |
| Symmetry imposed | C1 | C1 | C3 | C1 | C1 | C1 | C1 |
| Estimated accuracy of translations/rotations | 2.00/3.51 | - | 1.24/2.61 | 1.48/2.92 | 1.29/2.62 | 2.66/4.46 | 2.94/4.98 |
| Map sharpening B-factor | -158.6 | - | -83.8 | -90.4 | -87.6 | -85.3 | -75.2 |
| Unmasked Resolution at 0.5/0.143 FSC (Å) | 7.92/5.66 | 3.97/3.6 | 4.17/3.47 | 4.45/3.77 | 4.21/3.57 | 8.80/4.89 | 10.42/8.08 |
| Masked resolution at 0.5/0.143 FSC (Å) | 6.49/4.50 | 3.48/3.1 | 3.70/3.14 | 3.84/3.33 | 3.70/3.21 | 6.94/4.00 | 8.08/4.55 |
| **Model refinement and validation statistics** | | | | | | | |
| PDB | | | | | | | |
| Composition | | | | | | | |
| Amino acids | 3345 | 3283 | 3225 | 3274 | 3274 | 3274 | 3274 |
| Glycans | 57 | 57 | 57 | 57 | 57 | 57 | 57 |
| RMSD bonds (Å) | 0.014 | 0.015 | 0.012 | 0.013 | 0.018 | 0.014 | 0.016 |
| RMSD angles (◦) | 1.85 | 1.98 | 1.96 | 2.13 | 1.97 | 1.92 | 2.09 |
| Mean B-factors | | | | | | | |
| Amino acids | 59 | 61 | 84 | 116 | 90 | 64 | 66 |
| Glycans | 94 | 88 | 194 | 187 | 200 | 88 | 88 |
| Ramachandran | | | | | | | |
| Favored (%) | 91.00 | 92.73 | 92.74 | 93.70 | 92.9 | 91.28 | 91.45 |
| Allowed(%) | 8.19 | 6.87 | 6.82 | 5.86 | 6.55 | 8.19 | 7.83 |
| Outliers(%) | 0.81 | 0.40 | 0.44 | 0.43 | 0.56 | 0.53 | 0.72 |
| Rotamer outliers (%) | 2.23 | 1.01 | 3.75 | 1.76 | 3.11 | 1.97 | 2.22 |
| Clash score | 6.01 | 2.46 | 5.70 | 7.86 | 6.64 | 4.12 | 5.01 |
| C-beta outliers (%) | 0.29 | 0.65 | 0.46 | 0.52 | 0.16 | 0.49 | 0.89 |
| CaBLAM outliers (%) | 3.49 | 3.02 | 3.47 | 2.35 | 3.13 | 2.88 | 2.97 |
| CC (mask) | 0.63 | 0.78 | 0.85 | 0.87 | 0.86 | 0.64 | 0.59 |
| CC (volume) | 0.63 | 0.78 | 0.85 | 0.87 | 0.85 | 0.64 | 0.57 |
| MolProbity score | 2.11 | 1.49 | 2.21 | 2.03 | 2.19 | 1.93 | 2.03 |
| EMRinger score | 1.03 | 3.58 | 2.99 | 2.78 | 3.43 | 1.50 | 1.33 |
